# Inferred Structure of a Neuronal Circuit for Economic Decisions

**DOI:** 10.64898/2026.08.13.744684

**Authors:** Yuma Kanazawa, Kaining Zhang, Timothy G. Crimmins, Mahsa Khoshkhou, Heide Schoknecht, Gaia Tavoni, Camillo Padoa-Schioppa

## Abstract

Previous work suggests that different groups of neurons in orbitofrontal cortex (OFC) constitute the building blocks of a circuit in which economic decisions are formed. Here we used network inference analysis (Ising model) to shed light on the internal organization of this circuit. We examined populations of neurons recorded simultaneously, and inferred the functional couplings. We then computed a reduced, effective network (*EN*) where each node corresponded to an encoded variable. The *EN* had a recognizable structure, with enhanced couplings between input and output neurons supporting the same decision, and enhanced couplings between neurons encoding value variables with the same sign. This structure was highly reproducible across individuals and hemispheres. Importantly, it depended on the internal state of the animal and the behavioral conditions. The *EN* reproducibility decreased with the distance between cells but it increased with the number of cell pairs, suggesting that OFC operates as a single distributed assembly.

## Introduction

Choices between goods involve the representation and the comparison of subjective values. A broad literature based on clinical data^1–3^, functional imaging^4, 5^, lesions^6–8^, inactivation^9–12^, electrical stimulation^13, 14^, and neurophysiology^15–18^ links these operations to OFC. In particular, neurophysiology studies where monkeys chose between different juice flavors found that neurons in this area represent the offer values, the binary choice outcome, and the chosen value.^15, 18^ Notably, these variables capture both the input (offer values) and the output (chosen juice, chosen value) of the decision process, suggesting that the cell groups identified in OFC might constitute the building blocks of a circuit in which economic decisions are formed.^19^

Supporting this notion, computational models found that neural networks whose nodes match the cell groups found in OFC can generate binary decisions.^20–26^ However, current models leave open numerous questions. Most importantly, the structure of proposed networks is not based on empirical evidence, and different computational models present vastly different structures. In other words, the organization of the decision circuit, and thus the mechanisms governing economic decisions, remain poorly understood. To gain empirical insights on these fundamental questions, we recorded from neuronal populations in OFC while rhesus monkeys performed a juice choice task.^15^ We then used model-based inference^27–34^ to reconstruct the connectivity of the decision circuit.

Model-based inference estimates the functional couplings between pairs of neurons by taking into consideration not only the activity of the two cells (as in standard pairwise correlation), but also the interactions of these cells with other neurons recorded at the same time. Thus, it disentangles direct correlations between cells from indirect effects mediated by other recorded neurons. In particular, we fitted the spiking activity of neuronal populations with an Ising model.^35, 36^ We chose this over other models that support network inference^30, 37, 38^ for several reasons. First, of all the models restricted to pairwise couplings (i.e., excluding higher-order interactions), the Ising model is the least constrained, and thus the most general (maximum entropy).^35, 39, 40^ Second, the Ising model is generative, in the sense that it can predict higher-order correlations and collective activity patterns, even though it is fitted only to firing rates and pairwise interactions.^31, 34, 36, 41^ Third, previous work showed that Ising networks capture fundamental properties of circuit organization,^34, 42^ network-level coding,^43^ and memory consolidation emerging from Hebbian plasticity.^34^ Finally, the Ising model can be readily simulated,^33, 44^ which provides the opportunity to investigate the relationship between network structure and function. Of note, the Ising model is static in that it captures instantaneous co-activation patterns but not the temporal evolution of neuronal activity. Consequently, inferred couplings are symmetric (see **Discussion**). Importantly, inferred couplings capture statistical relations between cell pairs and should not be interpreted as faithful measures of synaptic efficacy.^30^

Here, we inferred an Ising network for each recording session.^36, 45^ Then, using a mean-field approach and pooling networks inferred in different sessions, we computed a reduced, effective network (*EN*), where nodes corresponded to signed variables encoded in OFC. For each pair of variables, we also computed the mean coupling strength (*MCS*), and we showed that *EN* and *MCS* are closely related. The *MCS* had a recognizable structure, with enhanced couplings between nodes supporting the same decision, reduced couplings between nodes supporting opposite decisions, and enhanced couplings between cells encoding variables offer value and chosen value with the same sign. This structure and the structure of the *EN* were highly reproducible across 3 monkeys and both hemispheres. Importantly, the *MCS* was modulated by the internal state of the animal and the behavioral conditions of the choice task. Its reproducibility decreased with the cell distance but it increased with the number of cell pairs, suggesting that neurons in OFC operate as a single assembly. Taken together, our results provide a first empirical assessment for the neural circuit supporting economic decisions.

## Results

Three male monkeys (D, E, F) performed a standard juice-choice task (**Fig.1AB**).^15^ In each session, animals chose between two juice flavors (A and B, with A preferred) offered in variable amounts. Offers were depicted as sets of colored squares on a monitor, and monkeys indicated their choice with a saccade. Choices consistently reflected a trade-off between juice flavor and juice quantity. For each session, choices were analyzed with a logistic regression, which provided measures for the relative value of the juices (*ρ*) and for the choice accuracy (*η*; **Fig.1C**).

**Figure 1.**
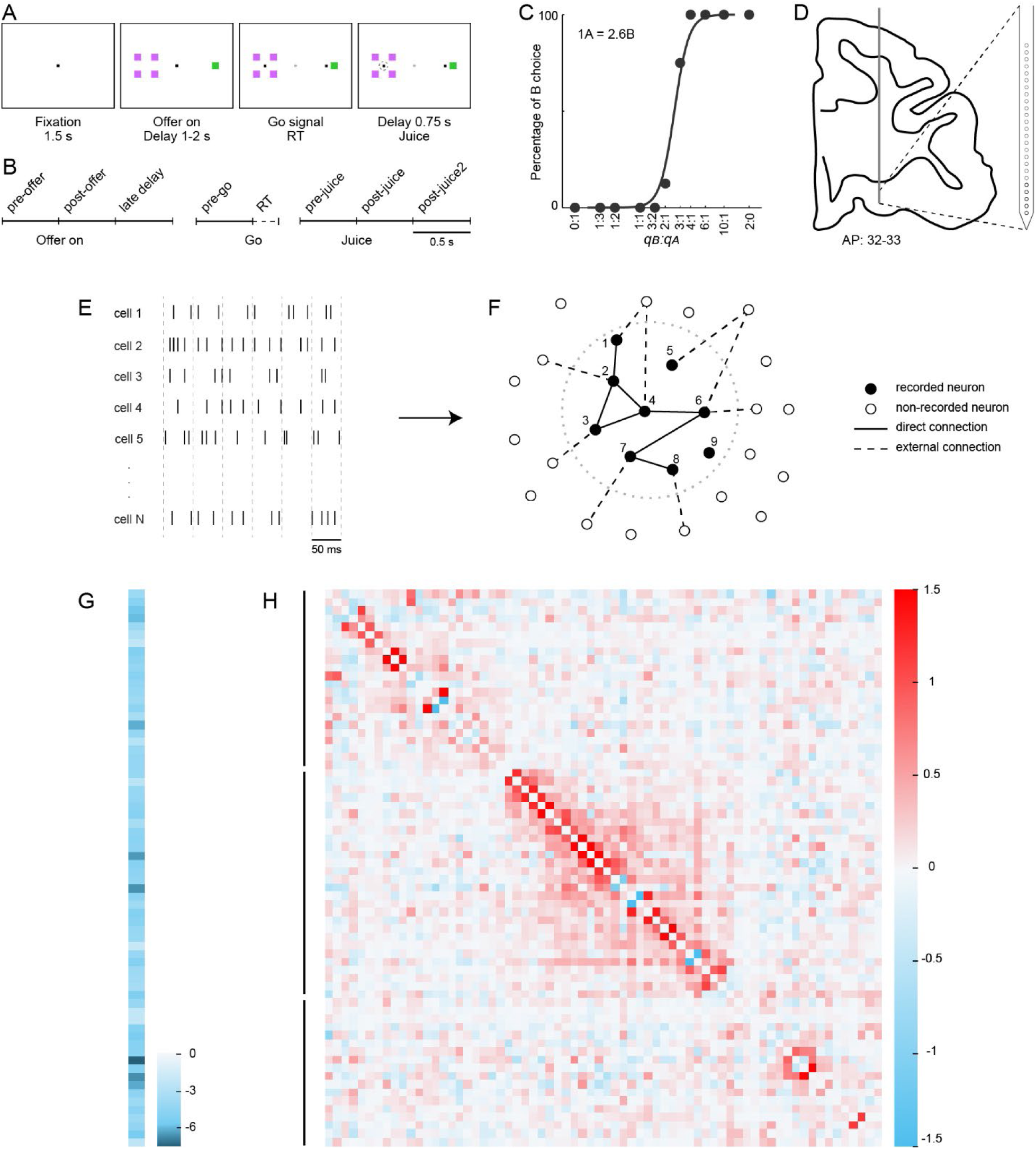
Choice task and network inference. **A**. Choice task. Each trial began with the animal gazing at a central fixation point. After 1.5 s, two offers appeared. Each offer was represented by a set of squares; the color and the number of squares indicated the juice flavor and the quantity, respectively. After a randomly variable delay (1-2 s), two saccade targets appeared by the offers (go signal). The animal indicated its choice by gazing at the corresponding target. After an additional 0.75 s delay, the chosen juice was delivered. **B**. Time windows for neuronal analysis. **C**. Choice pattern in one example session. The percentage of B choices (y-axis) is plotted against the quantity ratio *qB:qA* (x-axis, log scale; forced choices are plotted separately). The sigmoid curve was derived from a logistic regression. **D.** Schematic representation of the targeted recording area. The panel illustrates a coronal section and the vertical trajectory of the linear probe. **EF**. Network inference with the Ising model (cartoon). Spike times were parsed in 5-ms bins. The neuronal population activity in each time bin was represented as a vector ***σ*,** where each entry represented the state of a neuron (0 if silent, 1 if active). The inferred model was parametrized with inputs ℎ and couplings *J*. In panel F, filled circles represent recorded cells, empty circles represent non-recorded cells, solid lines represent internal connections (i.e., connections between pairs of recorded cells), and dashed lines represent external connections (i.e., connections between a recorded cell and a non-recorded cells). Connections are represented as unsigned and non-directional. Several motifs can be observed: cells 1 and 2 have a direct connection; cells 1 and 3 have no direct connection, but they have an internal indirect connection through cell 2; cells 5 and 6 have an external indirect connection; private inputs are represented for cells 2, 3, 6, 7, and 8. Inputs ℎ capture intrinsic cell properties and private inputs; couplings *J* capture direct connections and external indirect connections, but are not affected by internal indirect connections. **GH**. Inferred network, example session. This session included N = 80 neurons from 3 probes. Black bars on the left of panel H indicate the probe from which neurons were recorded. Panel G illustrates the external inputs (ℎ); panel H illustrates the pairwise couplings (*J*). Entries are color coded (see color bars). In panel G, most inputs were <0, capturing the fact that firing rates were generally low. In panel H, couplings could be either <0 or >0, with a bias towards the latter. The absolute strength of the couplings generally decreased with the distance between cells (distance from the diagonal), and was generally larger for neurons recorded with the same probe.

Extracellular recordings were performed in OFC in 85 sessions using multiple linear probes (**Fig.1D**). In any given session, we recorded 19-116 individual cells (median = 51 cells). The entire data set included 4,784 cells, and 149,025 cell pairs (**Suppl. Table 1; Suppl. Table 2; Suppl. Fig.1**). Unless otherwise noted, we only analyzed pairs of cells recorded simultaneously. Building on previous results,^19^ we classified individual neurons as encoding one of the variables characteristics of OFC, namely *offer value A*, *offer value B*, *chosen juice*, and *chosen value*. Each variable could be encoded with a positive or negative slope sign.^46^ Neurons that could not be assigned to any variable based on this analysis were labeled *unclassified* (**Methods, Suppl.** Table 3; Suppl. Table 4).

To quantify the functional connectivity, we examined spikes in 5-ms bins, and we fitted an Ising model to the mean firing probabilities and pairwise correlations of neurons recorded in each session (**Fig.1EF**; **Methods**). The Ising model is the least constrained model reproducing these low-order spiking statistics. Fitted parameters are a vector of inputs ℎ and a matrix of couplings *J*. For each cell *i*, input ℎ*_i_* captures the cell’s intrinsic tendency to fire as well as private inputs from other, unrecorded neurons. For each pair of cells *i* and *j*, coupling *J_ij_* represents a functional connection. A non-zero coupling *J_ij_* can emerge from direct anatomical connections between cells *i* and *j* and/or from indirect external connections through other, non-recorded cells. In contrast, indirect internal connections through other recorded neurons do not contribute to *J_ij_* .

**Fig.1GH** illustrates the network inferred for one example session where we used 3 probes and recorded from 80 cells. Inputs ℎ were generally negative, reflecting the fact that firing rates in OFC are typically low. Conversely, couplings *J* could be either positive or negative. On average, couplings were positive, and their absolute values tended to decrease with the distance between cells (**Suppl. Fig.2**; **Suppl. Fig.3)**.

**Figure 2.**
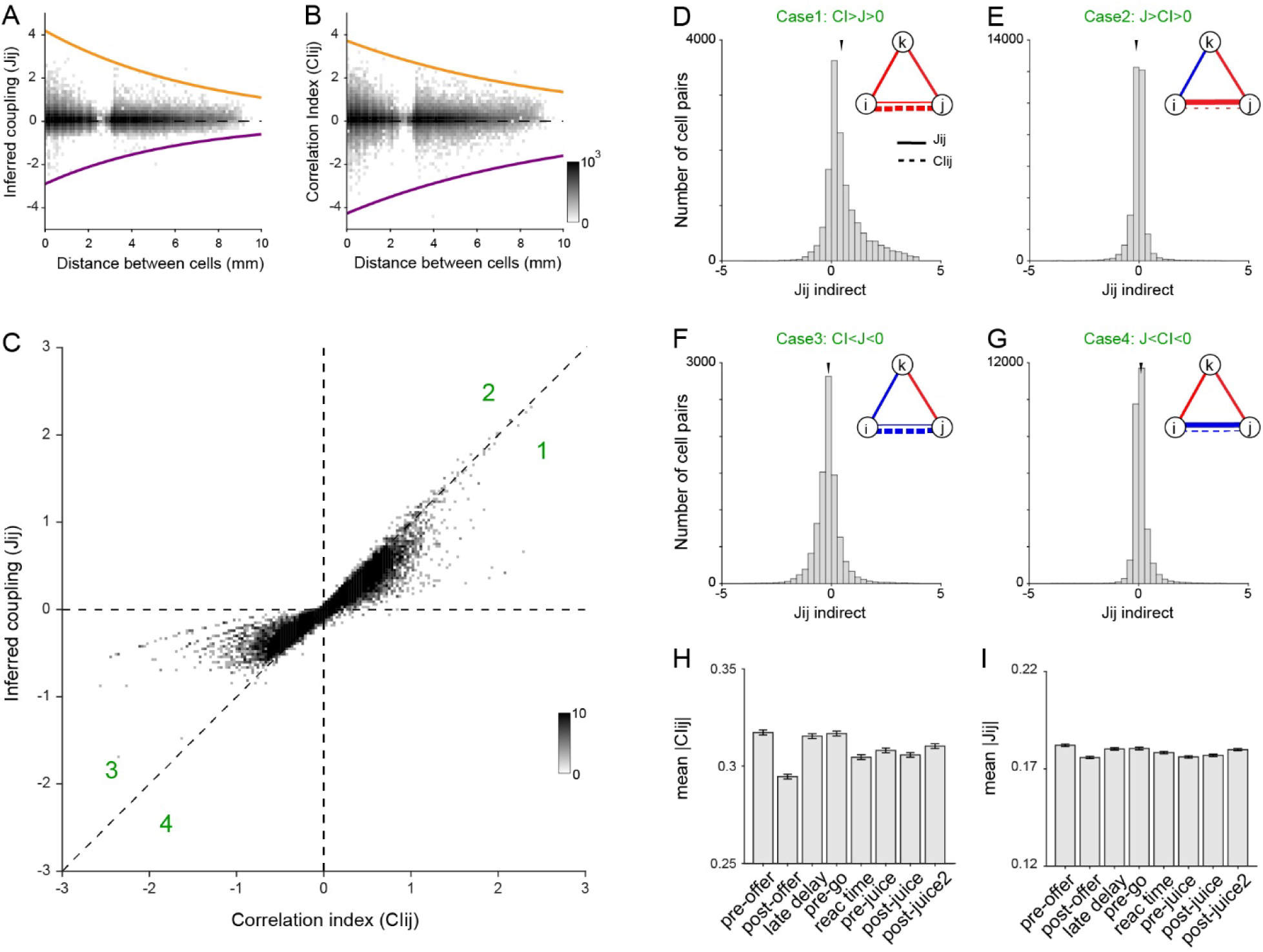
Inferred couplings and correlation indices. AB. Inferred couplings and correlation indices as a function of the distance between cells. Because of the large number of data points, here we display a pixel average (pixel size = 0.125 mm x 0.125). In both panels, orange and purple lines were obtained by dividing distances in 30 bins of equal size, by computing the maximum absolute value within each bin, and by fitting these maxima with an exponential function. Positive and negative values were analyzed separately. **C.** Pairwise comparison between inferred coupling (y-axis) and correlation index (x-axis). Here we display a pixel average (pixel size = 0.03 x 0.03). The two measures were highly correlated and nearly always (98% of pairs) had the same sign. However, on average, inferred couplings had smaller absolute values. **DEFG.** Differences between inferred couplings and correlation indices are explained by circuit motifs. We divided cell pairs in 4 groups depending on the relation between inferred coupling (*J*) and correlation index (*CI*), and their sign (see green numbers in panel C). For each group, we hypothesized a circuit motif that would explain the mean difference between *J* and *CI*. In each panel, the inset cartoon depicts the circuit motif for the corresponding case. In these cartoons, *i* and *j* are the two cells under consideration; *k* is a generic third cell recorded simultaneously; solid and dashed lines refer to inferred couplings and correlation indices, respectively; red and blue colors indicate positive and negative couplings or correlations, respectively; and the line weight indicates the strength of the coupling or correlation. For example, in case 1 (panel D), where *CIij* > *Jij* > 0, we hypothesized that the sum of indirect connections is positive. For each cell pair, we computed the indirect coupling, *Jij^indirect^*, as the sum of all 2^nd^ order indirect connections between cells *i* and *j* through other cells recorded simultaneously (**Eq.15**). The histogram in panel D illustrates the distribution of *Jij^indirect^* for case 1. Confirming predictions, the mean was significantly above zero (mean(*Jij^indirect^*) = 0.51; 2 10^-308^, two-tailed Wilcoxon Sign-rank test; N = 47,600). Panels E, F, and G illustrate the cartoon and the distribution of *Jij^indirect^* for cases 2, 3, and 4, respectively. In all cases, data were consistent with the hypothesized circuit motif (case 2: mean(*Jij^indirect^*) = −0.15; p = 3 × 10^-217^, case 3: mean(*Jij^indirect^*) = −0.037; p = 7 × 10^-12^, case 4: mean(*Jij^indirect^*) = 0.11; 3 10^-130^). Of note, the tails of the distributions of *J^indirect^* may deviate from these rules due to higher-order interaction effects mediated by longer chains of neurons. **HI**. Correlation indices and inferred couplings as a function of the behavioral time window. For panel I, the Ising network was inferred separately in each time window. Histograms illustrate absolute values (|*CIij*| and |*Jij*|). Confirming previous reports^50^, correlation indices differed significantly across time windows (p = 0.018, Kruskal-Wallis test). In contrast, inferred couplings did not differ significantly across time windows (p = 0.16, Kruskal-Wallis test).

**Figure 3.**
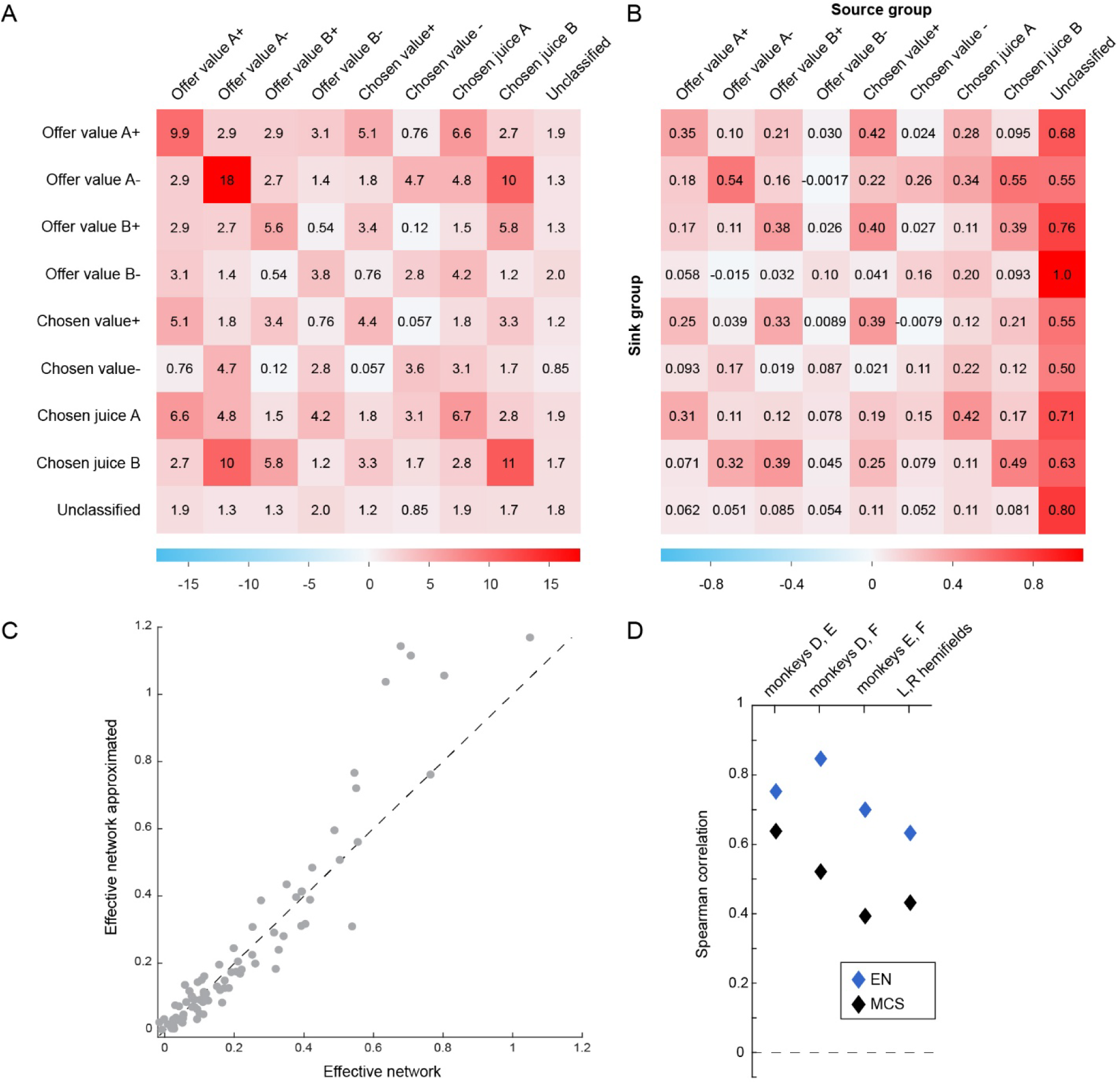
Reproducibility of the effective network. **A.** Mean coupling strength (*MCS*). Each entry is the weighted mean of inferred couplings between cells encoding the corresponding signed variables (**Methods**, **Eq.16**). **B.** Effective network (*EN*). Columns and rows are sources and sinks, respectively. Each entry *EN_uv_* captures the effective coupling from cell group v to cell group u, quantifying how activity in group *v* contributes to the activation of group *u* (**Methods**, **Eq.24**). **C.** Relation between *MCS* and *EN*. The x- and y-axis are the left- and right-hand side of **Eq.27**, respectively, and each data point represents one entry of *EN* (a pair of variables). The two measures are highly correlated (Spearman correlation = 0.95) and nearly indistinguishable for most entries. Substantial discrepancies are only observed for entries where *unclassified* cells are source. **D.** Reproducibility of the effective network. We computed *EN* and *MCS* for each of the 3 animals (D, E, F), and we examined their correlation across animals, pairwise. Similarly, we computed *EN* and *MCS* separately for the left (L) and right (R) hemispheres and then examined their correlation. The dashed line (correlation = 0) indicates chance level. Note that the lower correlations observed across hemispheres and pairs including monkey F are likely because *EN* and *MCS* were computed using fewer cell pairs (see **Suppl. Table 1,2**).

To assess the reliability of inferred couplings, we conducted a cross-validation analysis. For each session, we divided time bins into two sets depending on the bin number (even or odd), and we inferred a network for each set. Couplings inferred for the two sets were highly correlated (corr = 0.90 for statistically significant couplings) and their normalized difference was typically small (**Suppl. Fig.4AB**). Along similar lines, for each session, we inferred a network separately on easy and hard decision trials (see **Methods**). Couplings inferred for the two sets of trials were highly correlated (corr = 0.80 for statistically significant couplings; **Suppl. Fig.4CD**). Thus, inferred couplings were generally reliable and robust.

**Figure 4.**
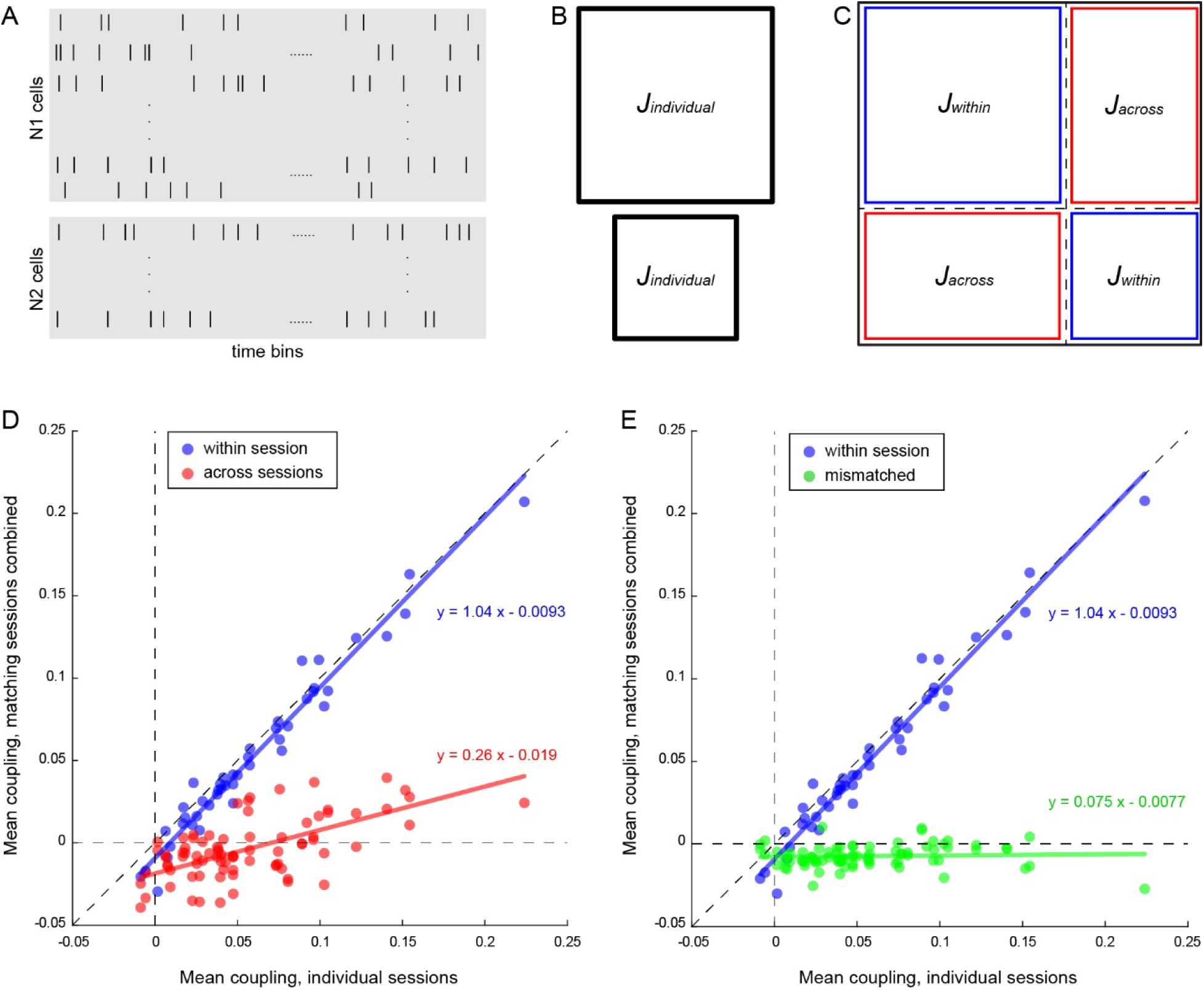
**Mean couplings are primarily driven by connections within the decision circuit. ABC**. Analysis of matching sessions. We identified pairs of matching sessions and matched trials across the two sessions (panel A). Next, we inferred the Ising network for individual sessions (*J^individual^* ; panel B) and for the two sessions combined (panel C). The network inferred for combined sessions was divided into 4 sectors – two on the diagonal with couplings between cells recorded simultaneously (*J**^ww^**^it^*^ℎ*in*^, blue), and two on the anti-diagonal with couplings between cells recorded in different sessions (*J^across^*, red). We conducted this analysis for all pairs of matching sessions and computed the average pairwise strength for each of the three treatments (*individual*, *within session*, and *across sessions*). **D.** Single coupling weakly depends on trial variables. The x-axis is the mean coupling obtained for individual sessions, the y-axis is the average coupling obtained for matching sessions, and each data point represents average pairwise strength between different pair of cell types. The two sets of data points represent mean couplings obtained *within session* (blue) and *across sessions* (red). Solid lines were obtained from Deming regressions. The 95% confidence intervals for the regression slopes are [0.97,1.14] (blue) and [0.19, 0.35] (red). **E.** Context variables alone do not induce functional couplings. We conducted the same analysis while randomly mismatching trials and task events (*mismatched* treatment). The 95% confidence intervals for the regression slopes are [0.98,1.15] (blue; *within session*) and [-0.039, 0.067] (green; *mismatched*). Confidence intervals for regression slopes were obtained from bootstrap tests (N=50,000).

Importantly, the Ising model makes it possible to predict the probability that any given neuron emits a spike in any time bin, conditioned on whether other cells recorded simultaneously spike in the same time bin. Although predicting spikes was not the main focus of this study, we conducted analyses to verify the validity of these predictions. We found that spiking predictions obtained from inferred networks were significantly more accurate than those derived from mean-field models based on firing rates averaged across trials or across cells (see **Methods**, **Suppl. Fig.5**).

**Figure 5.**
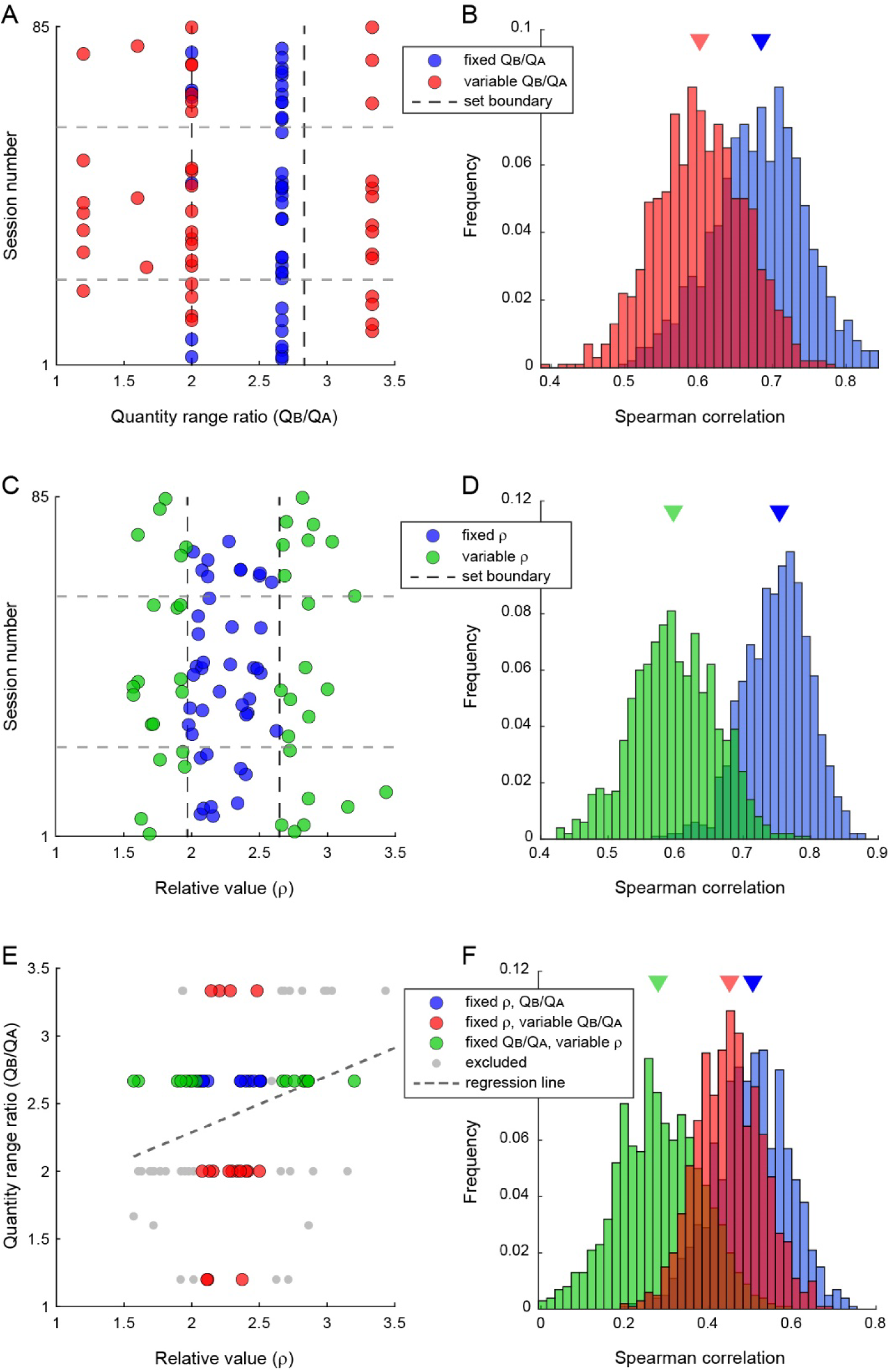
The internal state of the animal modulates anatomical connections. **A**. Distribution of quantity range ratio (*QB*/*QA*; N = 85 sessions). The x-axis and y-axis represent, respectively, the quantity range ratio and the session number (ordered chronologically pooling data from the 3 monkeys). Sessions were split into two sets based on the distance from the mean of the distribution (near-center, N = 42 sessions, blue color; far-from-center, N = 43, red color). Vertical dashed lines indicate the split boundaries. Sessions at the boundary (*QB*/*QA* = 2) were assigned randomly to one set or the other in proportions such that the two sets would have roughly equal number of sessions. Horizontal dashed lines separate data points from the three animals. **B**. Split-half reproducibility of the *MCS* matrix. For each set of sessions, we pooled cell pairs across sessions, randomly divided them in two pools, computed the two *MCS* matrices, and calculated their Spearman correlation. We repeated these operations 1,000 times and obtained a distribution of Spearman correlations. The distributions obtained for the two sets of sessions were significantly different (median Spearman correlations = 0.69 (blue) and 0.60 (red); rank-biserial r = 0.66; p = 1 × 10^-143,^ Mann-Whitney U test). Triangles indicate distribution medians. In other words, the quantity range ratio *QB/QA* significantly affected the structure of the *MCS* network. **CD**. Mean coupling reproducibility and relative value (*ρ*). Same analysis as in panels AB conducted splitting sessions according to the relative value *ρ*. The distributions of Spearman correlations obtained for near-center sessions (blue) and far-from-center sessions (green) were significantly different (median Spearman correlations 0.75 (blue) and 0.61 (green); rank-biserial r = 0.93; p = 3 × 10^-285,^ Mann-Whitney U test). In other words, the structure of the mean coupling was significantly affected by the internal state of the animal. Furthermore, the size of this effect was substantially larger than that measured in panel B. **E**. Joint distribution of quantity range ratios and relative values. The two measures were significantly correlated (Pearson’s r = 0.31, p = 0.0044). The dashed line was derived from a linear regression. Sessions were divided into three sets using a conditioned median-split: center (blue, N = 17; |z| < 0.6 standard deviation on both axes), far-*ρ* (green, N = 17; dissimilar *ρ*, conditioned on similar *QB*/*QA*), and far-*QB*/*QA* (red, N = 17; dissimilar *QB*/*QA*, conditioned on similar *ρ*). Grey points indicate other, excluded sessions (N = 34). **F**. Split-half reproducibility of the mean coupling strength. The distribution of Spearman correlations for each set was obtained as in panel B. The three distributions were all significantly different from each other (center vs. far-*ρ*: rank-biserial r = 0.91, p = 3 × 10⁻^269^; center vs. far-*QB*/*QA* : rank-biserial r = 0.32, p = 5 × 10⁻^36^; far-*QB*/*QA* vs. far-*ρ*, rank-biserial r = 0.81, p = 3 × 10⁻^220^; all p values from Mann-Whitney U tests).

### Inferred couplings and correlation indices

Historically, pairwise relations between cells recorded simultaneously have been studied primarily through noise correlation.^47–50^ A closely related metric, the correlation index, is directly comparable to Ising couplings^36^ and will be discussed here. Given two cells *i* and *j*, the correlation index is defined as *CI_ij_* = log(*p_ij_*⁄(*p_i_ p_j_*)), where *p_i_* is the probability that cell *i* spikes in a 5-ms bin, and *p_ij_* is the probability that both cells *i* and *j* spike in a 5-ms bin. The fundamental difference between correlation indices and Ising couplings is that the former are computed based on the activity of two cells alone; in contrast, the latter are computed taking into consideration the activity of all other neurons recorded simultaneously. Notably, because they are computed all at once, Ising couplings provide a coherent model. We investigated the relation between Ising couplings and correlation indices.

Both correlation indices and inferred couplings could be positive or negative, and both of them decreased in absolute value as a function of the distance between cells (**Fig.2AB**). Over the population, the two measures were highly correlated (Pearson’s r = 0.92, **Fig.2C**). However, absolute values |*CI_ij_*| and |*J_ij_* | differed significantly (p = 2 × 10^-308^, signed-rank test). We hypothesized that this difference reflected functional connections with other cells in the network. To appreciate why that might be, consider two neurons *i* and *j* and assume that both *CI_ij_* and *J_ij_* are positive. As noted, *CI_ij_* is computed based on the activity of the two cells alone, while *J_ij_* also takes into consideration all other cells recorded simultaneously. If a third cell *k* in the network has positive functional connections with both *i* and *j*, these connections will increase the correlation index *CI_ij_*, but they will not affect the coupling *J_ij_* . Similarly, if a third cell *k* has a positive functional connection with one cell and a negative functional connection with the other, these connections will reduce *CI_ij_* but not affect *J_ij_* . More generally, functional connections with other cells in the recorded network will induce differences between *CI_ij_* and *J_ij_* .

To test our hypothesis, we defined, for any two cells *i* and *j*, the (internal) indirect coupling *J^indirect^*, which captured the interaction between cells *i* and *j* mediated by all the other neurons recorded simultaneously (see **Methods**). We then analyzed differences between *CI_ij_* and *J_ij_* as a function of *J^indirect^*. (Note: external indirect connections are not considered in this section.)

Referring to **Fig.2C**, we examined 4 cases. In case 1, *CI_ij_* > *J_ij_* > 0. For the reasons detailed above, we expected that indirect couplings would be generally positive. Indeed, we found that mean(*J_ij_^indirect^*) was significantly larger than zero (mean(*J_ij_^indirect^*) > 0; p = 2 × 10^−308^, Wilcoxon Sign-rank test **Fig.2D**). In case 2, *J_ij_* > *CI_ij_* > 0. As discussed above, we expected indirect couplings to be generally negative, and this is indeed what we found (mean(*J^indirect^*) < 0; p = 3 × 10^-217^, Wilcoxon Sign-rank test; **Fig.2E**). In case 3, *CI_ij_* < *J_ij_* < 0. We expected indirect couplings to be negative, and our analysis confirmed this prediction (mean(*J^indirect^*) < 0; p = 7 × 10^-12^, Wilcoxon Sign-rank test; **Fig.2F**). Finally, in case 4, *J_ij_* < *CI_ij_* < 0. We expected indirect couplings to be positive, and our analyses confirmed the prediction (mean(*J^indirect^*) > 0; p = 3 × 10^-130^, Wilcoxon Sign-rank test; **Fig.2G**).

Previous work found that noise correlations vary over the course of a trial, and drop specifically at the time when perceptual or value-based decisions are formed.^50, 51^ In attentional tasks, this drop is strongly linked to the behavioral performance.^51^ The short time scale on which it occurs suggests that the correlation drop is driven by top down input, as opposed to local changes in synaptic strength. As noted above, inferred couplings capture statistical interactions, not anatomical properties. At the same time, because they are computed net of indirect interactions through other recorded cells, inferred couplings are presumably closer to actual synaptic strengths compared to noise correlations or correlations indices. This consideration suggests that modulation over the course of a trial should be substantially reduced for inferred couplings. Our analyses validated this prediction. First, we confirmed that correlation indices dropped significantly during the post-offer time window (p = 0.018, Kruskal-Wallis test; **Fig.2H**), as previously observed for noise correlations.^50^ In contrast, the drop measured for inferred couplings in the same time window was much more subtle and did not reach statistical significance (p = 0.16, Kruskal-Wallis test; **Fig.2I**).

### Mean coupling strength and effective network

The primary goal of this study was to gain insights into the mechanisms underlying economic decisions. Large connectivity matrices where entries are single neurons (**Fig.1H**) are hard to interpret as a mechanism. In contrast, we sought to condense inferred networks to an interpretable form, and to study the decision circuit at the mesoscale.

To do so, we first computed a 9×9 matrix of mean connection strengths (*MCS*). In this matrix, each row and each column corresponded to a signed variable (i.e., a cell group). Given cell groups *u* and *v*, entry *MCS_uv_* was defined as the weighted average of all inferred couplings between cells encoding variables *u* and *v* available in our data set (**Fig.3A**). Of note, *MCS* was computed correcting for the fact that the strength of inferred coupling decreased with the number of cells recorded simultaneously (**Suppl. Fig.6**; **Methods**).

**Figure 6.**
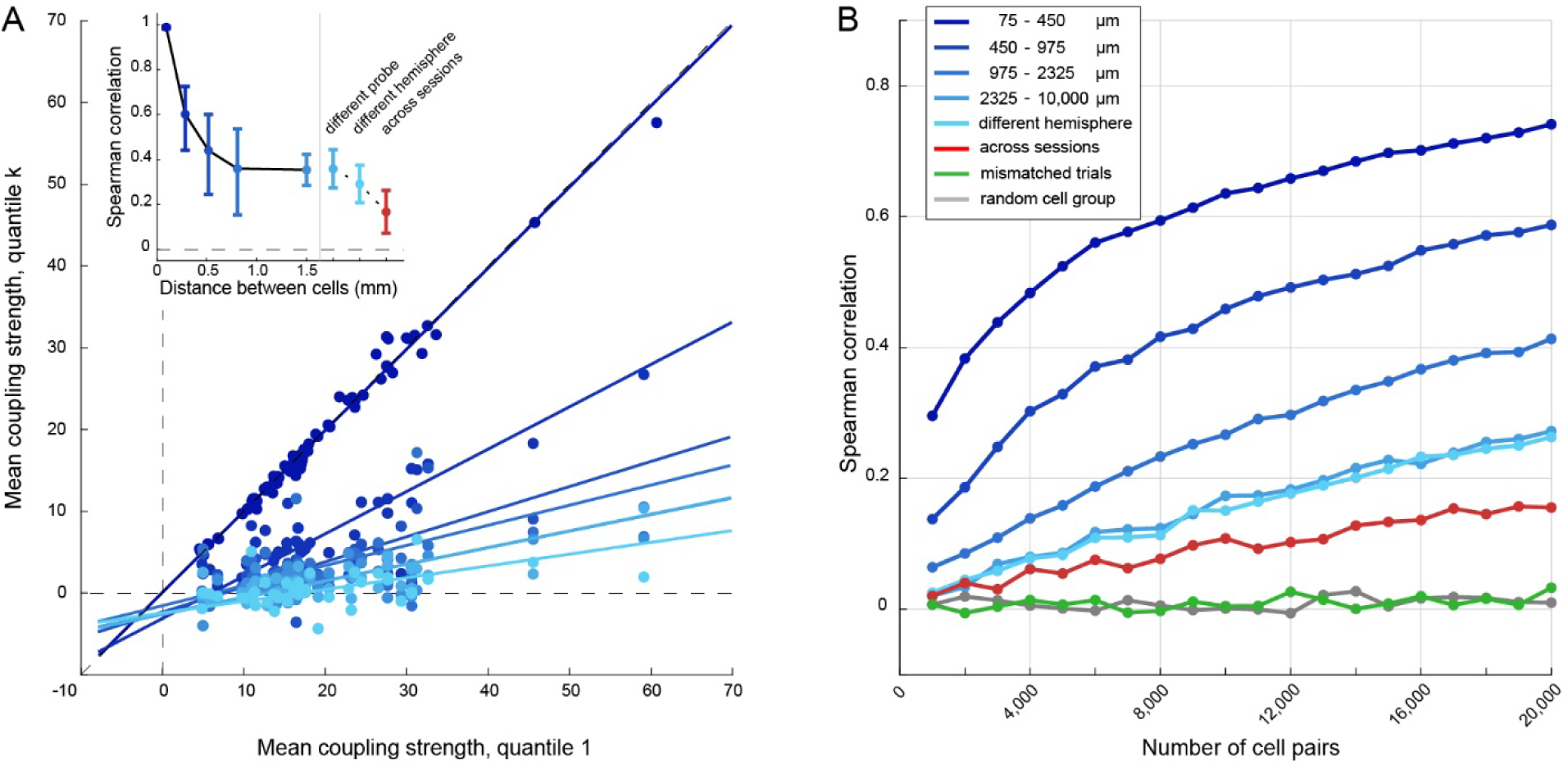
Effective network, cell distance, and sample size. **A.** Structure of *MCS* as a function of the cell distance. Cell pairs were divided into 7 pools by distance (see **Methods**). For each pool of cell pairs, we computed the *MCS* matrix. In the main plot, the x-axis is the coupling strength for pool 1 (shortest distance), the y-axis is the coupling strength for pool k = 1…7, and each data point represents one entry of *MCS* (N = 45 data points per pool). Different shades of blue represent different pools (see inset), and solid lines are from Deming regressions. For k = 1, data points were derived from a split-half procedure (**Methods**). In the inset plot, the x-axis is the mean distance between cell pairs in a pool, the y-axis is the Spearman correlation between pool k and pool 1, and each data point represents one pool (same colors as in the main plot). The red data point represents the *across sessions* treatment. For the 3 pools at distances >1 mm and the *across sessions* treatment, the correlation was estimated with a bootstrap procedure (see **Methods**). **B.** Structure of *EN* as a function of the number of cell pairs. The reproducibility of *EN* (Spearman correlation, y-axis), is plotted against the number of cell pairs (x-axis), for different cell distances (shades of blue; see legend). The red line was computed using couplings from the *across sessions* treatment. The green line was computed using inferred couplings from the *mismatched* treatment. The gray line was computed by randomizing cell identities before computing the *EN*.

Second, using a mean-field approach, we derived a coarse-grained 9×9 effective network (*EN*; **Fig.3B**) where each entry *EN_uv_* quantified the increase in mean activity for cell group *u* (sink) conditioned on the increase in mean activity for cell group *v* (source). In other words, *EN* captured the effective statistical influence of each cell group onto other cell groups. Notably, the matrix *EN* was not symmetric (see **Methods**).

If the number of cells recorded simultaneously is sufficiently large, entries in *EN* can be approximated by the product *EN_uv_* ≈ *P_v_ MCS_uv_*, where *P_v_* is the probability that a cell in OFC encodes variable *v* in a session (see **Methods, Suppl. Table 3**). As illustrated in **Fig.3C**, this condition held in our data, in the sense that the two matrices *EN* and *P* ∗ *MCS* where very similar (Spearman correlation = 0.95). In the remainder of this study, we analyzed the structure and properties of matrices *MCS* and *EN*.

### Structure and reproducibility of the effective network

Several aspects of the *MCS* matrix were most noteworthy (see highlights in **Suppl. Fig.7**). First, couplings between classified and unclassified cells were generally weak and seemed rather unstructured. Their mean value could be taken as a baseline comparison for other entries of the *MCS* matrix. Second, for each of the 4 variables (*offer value A*, *offer value B*, *chosen value*, *chosen juice*), couplings between cells encoding it with the same sign were enhanced, while couplings between cells encoding it with opposite signs were at or below baseline. Third, couplings between offer value cells and chosen juice cells were enhanced when the two neurons supported the same decision (e.g., *offer value A+* ↔ *chosen juice A*; *offer value A–* ↔ *chosen juice B*) compared to when they supported opposite decisions (e.g., *offer value A+* ↔ *chosen juice B*). These patterns resonate with the structure of current computational models.^22, 52^ Fourth, couplings between *offer value* cells associated with different juices were unstructured and barely above baseline. Fifth, couplings between *offer value* cells and *chosen value* cells were enhanced when the encoding sign was the same (+ or –) and depressed when the encoding sign was opposite. While we cannot assess the sign of the encoding for *chosen juice* cells with the current task (but see [^46^]), this observation suggests the presence of two, relatively disconnected networks of positively and negatively tuned cells.

Many of the same traits could also be observed in the *EN* (**Fig.3B**). The main difference – i.e., that connections originating from *unclassified* cells were enhanced in *EN* – was due to the large size of this cell group.

We aimed to assess whether the structure of *EN* and the structure of *MCS* were reproducible. Thus, we computed *EN* and *MCS* separately for each of the 3 animals participating in the experiment (**Suppl. Fig.8**). We then quantified the pairwise similarity between matrices. Both *MCS* and *EN* were highly reproducible (mean Spearman correlation >0.5 and >0.8, respectively; **Fig.3D**). Along similar lines, we computed *EN* and *MCS* using exclusively pairs of cells recorded in the left hemisphere and, separately, pairs of cells recorded in the right hemisphere. For both *EN* and *MCS* we found a strong correlation (Spearman correlation >0.4 and >0.7, respectively; **Fig.3D**). Thus, the structure of the decision circuit is reproduced in each hemisphere and preserved across individuals.

### Mean coupling strengths are primarily driven by anatomical connections

It is natural to regard the structure of *MCS* illustrated in **Fig.3A** as determined by direct or indirect anatomical connections between neurons that participate in the decision process. However, functional connections between pairs of cells, and thus the structure of *MCS*, might be partly or entirely driven by behavioral variables, which include event variables (the time of task events such as offer onset and juice delivery), trial variables (offer quantities *q_A_*, *q_B_* and the chosen juice), and context variables (quantity ranges *Q_A_*, *Q_B_* and the relative value *ρ*). In other words, even two cells that are not directly or indirectly connected (e.g., two cells recorded in different animals) could present nonzero functional couplings simply because they activate at the same time relative to behavioral events and/or because they encode correlated variables. We conducted a series of analyses to shed light on this issue.

We identified in our data set pairs of matching sessions – i.e., sessions where juice flavors, trial types, and relative values were nearly identical. For each pair of matching sessions, we matched trials such that offer quantities (*q_A_*, *q_B_*) and choices would be the same for the two sessions. We defined two time windows aligned with offer onset and juice delivery; we then concatenated the two time windows for each trial, and all trials for each session (**Methods**). Thus we obtained two data matrices for the two sessions. In each matrix, columns were time bins and rows were cells (**Fig.4A**). The two data matrices had the same number of columns (because times were matched), but not necessarily the same number of rows (because cell numbers were generally unequal).

First, we inferred the network for each session (*J^individual^*; **Fig.4B**). Then, we concatenated the two matrices as though all cells had been recorded in the same session, and we inferred the network for the combined pseudo-session. The resulting network was divided into 4 sectors (**Fig.4C**). Two sectors on the main diagonal included couplings between pairs of cells recorded simultaneously (*J****^within session^***). The other two sectors included couplings between pairs of cells recorded in different sessions (*J^across^ ^sessions^*). We reasoned that these functional couplings could not be driven by anatomical connections. Therefore, any structure in *J^across^ ^sessions^* couplings would necessarily be induced by some behavioral variable.

In another treatment, we repeated this analysis, but we randomly mismatched trials and misaligned the timing of behavioral events between the two matching sessions (**Methods**). In this treatment, functional couplings between pairs of cells recorded in different sessions were labeled *J^mismatc^*^ℎ*ed*^ . Since this procedure scrambled all event and trial variables, any structure in *J^mismatc^*^ℎ*ed*^ would necessarily reflect a contextual variable (i.e., *Q_A_*, *Q_B_*, or *ρ*).

We conducted these analyses for each pair of matching sessions (N = 78). Pooling the results, we computed average coupling for each of the 3 treatments (*within session*, *across sessions*, *mismatched*). Finally, we plotted the three sets of couplings against the corresponding couplings obtained from the standard analysis of individual sessions (**Fig.4DE**). Several aspects of the results were noteworthy. (1) As expected, *within session* couplings were nearly identical to *individual session* couplings; indeed, they captured the same interactions and differed only minimally because *within session* couplings were computed concurrently with *across sessions* couplings. (2) *Across sessions* couplings were substantially lower than *individual session* couplings. However, (3) *across sessions* couplings were generally nonzero, and (4) they were significantly correlated with *individual session* couplings (r = 0.45; p = 2 × 10^-5^; Spearman correlation). Conversely, (5) *mismatched* couplings were close to zero, and statistically independent from *individual session* couplings (r = 0.20; p = 0.07; Spearman correlation). These results were replicated for individual animals (**Suppl. Fig.8**) and in a variant of the analysis where each session was matched only once (not shown).

We interpret these observations as follows. First, inferred couplings *J* were primarily driven by anatomical connections (implied by 2). However, pairs of cells lacking direct or indirect connections presented functional couplings (implied by 3). These couplings were structured and not random (implied by 4). They depended on the functional properties of the two cells, and they were induced by behavioral variables that varied on each trial (*q_A_*, *q_B_*, the chosen juice, and the timing of behavioral events). Once these variables were controlled, effective couplings lost any structure (implied by 5).

### Connection strengths are modulated by the internal state and the behavioral context

The relative value *ρ* is the quantity ratio *q_B_*/*q_A_* that makes the animal indifferent between the two juices. It is a parameter that reflects the subjective preferences and the motivational state of the animal. For example, for any two juice flavors, *ρ* tends to increase when the animal becomes less thirsty. Conversely, quantity ranges *Q_A_* and *Q_B_* are set by the experimenters. The relevance of these variables to the decision process follows from the fact that value encoding cells in OFC undergo range adaptation.^53, 54^ Unless the decision mechanisms compensate it in some ways, range adaptation would induce maladaptive choices biases.^55, 56^ These considerations and theoretical work on optimal coding lead to the prediction that context variables *ρ*, *Q_A_*, and *Q_B_* should inform the decision circuit.^55, 56^ Importantly, **Fig.4E** indicates that context variables alone do not induce any functional coupling between pairs of cells in OFC. At the same time, context variables could shape the decision circuit by modulating the strength of anatomical connections. We set up to test this hypothesis.

First, we considered the distribution of quantity range ratios *Q_B_*/*Q_A_*, which varied roughly between 1 and 3.5 (**Fig.5A**). We divided the 85 sessions in two sets based on the median distance from the center of the distribution. The two sets had roughly the same number of sessions (42 and 43, respectively) and mean value of *Q_B_/Q_A_*, but they had different *Q_B_/Q_A_* dispersion. As a result of this procedure, we could examine the reproducibility of the *_MCS_* matrix in two conditions, with nearly fixed *Q_B_*/*Q_A_* (near-center sessions) and with highly variable *Q_B_*/*Q_A_* (far-from-center sessions). For each set of sessions, we examined the reproducibility of the *MCS* matrix using a bootstrap, split-half correlation procedure. We randomly divided the population of inferred couplings in two pools, computed two *MCS* matrices, and calculated their similarity (Spearman correlation). We repeated these operations N = 1,000 times, and we obtained a bootstrap distribution for the Spearman correlation. Finally, we compared the distributions obtained for the two sets of sessions. Inspection of **Fig.5B** reveals that the reproducibility of *MCS* was significantly higher when *Q_B_*/*Q_A_* was nearly fixed compared to when it was highly variable (rank-biserial r = 0.93; p = 4 × 10^-282^, Mann-Whitney test). In other words, the quantity range ratio *Q_B_*/*Q_A_* significantly affected the structure of the network.

We repeated this analysis dividing sessions according to the distribution of relative values *ρ*, which varied roughly between 1.5 and 3.5 (**Fig.5C**). Again, we divided sessions in two sets based on the median distance from the center of the distribution. For each set of sessions, we examined the reproducibility of *MCS* using a bootstrap, split-half correlation procedure. We found that the reproducibility of *MCS* was significantly higher when the relative value *ρ* was nearly fixed compared to when it was highly variable (rank-biserial r = 0.59; p = 5 × 10^-116^, Mann-Whitney test; **Fig.5D**). In other words, the internal state of the animal, captured by the relative value *ρ*, significantly shaped the structure of *MCS*. Notably, the size of this effect was roughly twice as large as that observed in **Fig.5B**.

We noted that variables *ρ* and *Q_B_/Q_A_* were significantly correlated across sessions in our data set. Consequently, the separation between histograms illustrated in **Fig.5B** could be partly induced by differences in *ρ*, as opposed to differences in *Q_B_/Q_A_* per se. However, a more granular analysis in which we effectively removed this correlation confirmed the results discussed above. The structure of *MCS* significantly depended on both the internal state (*ρ*) and the external conditions (*Q_B_/Q_A_*), and the effect of the internal state was substantially larger (**Fig.5EF**).

### Mean coupling strength, cell distance, and sample size

In a final set of analyses, we examined how *MCS* and *EN* varied as a function of the distance between cells and the number of cell pairs.

First, we noted that both the strength and the signal-to-noise ratio of mean couplings generally decreased as a function of the cell distance (**Suppl. Fig.3**). To examine how the structure of *MCS* varied with the cell distance, we considered the distribution of cell-pair distances (**Suppl. Fig.1**) and divided cell pairs in 7 pools based on the distance (pool 1 corresponding to the shortest distance; see **Methods**). For each distance pool k, we computed the *MCS* matrix, which we labeled *MCS*^(*k*)^. We then plotted entries of *MCS*^(*k*)^ against the corresponding entries of *MCS*^(1)^ (**Fig.6A**). We computed the Spearman correlation across entries, and we performed a linear Deming regression. These operations were repeated for each k = 1…7, and we examined how Spearman correlation and regression slope varied as a function of k. Remarkably, the two sets of *MCS* entries were correlated for all values of k. However, the Spearman correlation and the regression slope orderly decreased as a function of k. In other words, as the distance between cell pairs increased, the *absolute* strength of all connections decreased, but the *relative* strength of different connections (i.e., the structure of *MCS*) remained similar. The correlation appeared to reach a low plateau at distances ≥0.8 mm. Note, however, that this plateau was significantly above chance even at large distances, suggesting that even connections between distant cells maintained a structure similar to that observed in **Fig.3D**.

A different analysis revealed that our measures of *EN* reproducibility strongly depended on the number of cell pairs included in the calculation. In this case, we examined all cell pairs, and we divided the distribution of distances into 5 quantiles (∼20,000 cell pairs per quantile). For the first distance quantile, we computed two *EN*s based on two sets of N = 1,000 randomly selected cell pairs, and we calculated their similarity (Spearman correlation). To obtain a robust measure, we repeated this operation 100 times and noted the mean(Spearman correlation) = 0.18. We then repeated these operations computing *EN*s with increasingly larger numbers of cell pairs (up to N = 10,000), and for different quantiles of the distribution of distances (**Fig.6B**). As already noted, the reproducibility of *EN* decreased with the distance between cell pairs. Remarkably, however, even when the network was computed using pairs of cells at large distances (>2 mm), the *EN* reproducibility was substantially above chance, and it continued to increase as the number of cell pairs increased.

To assess whether this result was driven by behavioral variables, we repeated the analysis using *across sessions* couplings obtained from matching sessions (see above). As already noted in **Fig.4D**, these couplings were nonzero, and their reproducibility increased with the number of cell pairs. Critically, the *EN* reproducibility obtained for cell pairs at long distance (light blue) was substantially higher than that obtained for cell pairs recorded in different sessions (red), and this difference increased as a function of the number of cells pairs included in the analysis (**Fig.6B**).

For a control, we repeated this analysis after randomly reassigning each neuron to a signed variable before computing the effective network. As expected, the resulting *EN*s were completely uncorrelated. Analyses of *MCS* yielded similar results (not shown).

Taken alone, **Fig.6A** might suggest that the decision process is dominated by short-distance connections. However, in any cortical area, the number of cell pairs at distance *d* increases as a function of *d* (to a first approximation, it scales as *d^2^*). In this light, the results of **Fig.6B** suggest that OFC operates as a single decision circuit structured by both short- and long-distance connections.

## Discussion

Previous neurophysiology studies found that neurons in OFC represent the offer values (decision input), the chosen value, and the binary choice outcome (decision output). This observation and other results^8, 14, 16, 19^ suggest that OFC contains the building blocks of a decision circuit. A fundamental question concerns the organization of this neural circuit – i.e., how different groups of neurons in OFC are connected with each other and with other brain regions. Addressing this issue is challenging because standard neurophysiology techniques don’t provide measures of connectivity. Some insights into the circuit organization can come from the analysis of noise correlation, or the closely related *CI*. ^47, 50^ However, noise correlation and *CI* are computed between pairs of cells taken in isolation and thus confound direct interactions with indirect correlations mediated by other cells. The possibility to record large neuronal populations simultaneously allows more powerful, model-based approaches. In particular, here we inferred the functional connectivity in OFC using the Ising model.^29, 34–36^ Our results may be summarized as follows.

First, inferred couplings were reliable, robust, and stable across time windows. Although inferred couplings were strongly correlated with *CI*s, there were also systematic differences between the two measures. Specifically, the absolute value of the *CI* was typically larger than that of the corresponding inferred coupling. This difference was accounted for by indirect connections with other neurons recorded in the same session. Other analyses revealed that predictions of spike timing for individual cells based on inferred networks were significantly more accurate than those derived from average firing rates across trials or across cells. These results validate network inference analysis for this neural circuit.

Second, using a mean-field approach and pooling inferred networks from multiple sessions, we defined an *EN* where each node corresponds to a signed variable encoded in OFC. In essence, each entry in *EN* captures the activity increase of one group of cells conditioned on the activity increase of another group of cells. We also computed the closely related *MCS*, which captures the average coupling strength between cells encoding specific pairs of variables. Both *EN* and *MCS* had a recognizable structure. In particular, *MCS* presented enhanced couplings between input and output cells supporting the same decisions. Most remarkably, the structure of both *EN* and *MCS* was highly reproducible across monkeys and hemispheres. Since inferred couplings generally presented high variability, observing the reproducibility of *EN* and *MCS* required a substantial number of cell pairs (∼10^3^-10^4^). It is worth noting that, while large for neurophysiology standards, this number of cell pairs is modest in the perspective of a cortical brain area.

Third, choices between different goods are dictated by subjective preferences and by the internal state of the animal, both of which are captured by the relative value of the juices (*ρ*). Thus, the decision mechanisms should critically depend on *ρ*. Along similar lines, theoretical work proposed that the decision circuit adapts to the ranges of offer values – a necessary adjustment to avoid undue choice biases induced by range adaptation.^55, 56^ Consistent with these notions, we found that the structure of *MCS* was significantly shaped by the relative value of the juices and by the ranges of offered values. Importantly, these variables were not encoded by individual neurons per se, and they did not couple the spiking activity of cell pairs in OFC directly. Instead, these variables shaped the decision circuit by modulating the strength of anatomical connections.

Fourth, the absolute value of inferred couplings and the associated SNR decreased as a function of the distance between cell pairs. Consequently, for any given number of cell pairs, the reproducibility of *EN* decreased as a function of distance. However, the reproducibility remained above chance even for large distances and increased with the number of cell pairs. In other words, connections between pairs of cells at large distances or in different hemispheres were weaker on average, but similar in structure to those between cell pairs at short distances. This observation resonates with studies showing that a large fraction of connections in cortex are non-local.^57–59^ Considering that the number of cell pairs increases as a function of the distance, our observations suggest that neurons in OFC operate as a single, distributed circuit.

One important question is whether the emerging network is a credible computational model to generate binary decisions. We address this issue in a related study, where we show that the inferred networks indeed support attractor transitions consistent with choice.^44^

These results should be considered in the light of several caveats. First, our analysis assumed that the representation of decision variables in OFC is categorical – i.e., that each neuron encoded one variable (or no variable, for unclassified cells). However, recent analyses suggest that the representation in OFC might be more mixed than initially thought.^60, 61^ Thus, future work should assess the network connectivity while relaxing the assumption of categorical encoding. Second, computational studies strongly suggest that the competitive winner-take-all process underlying economic decisions is shaped by inhibition.^22, 26, 62–67^ We cannot firmly distinguish between excitatory (pyramidal) cells and inhibitory interneurons in our data set. Inferred Ising couplings are signed and, indeed, ∼44% of them were <0 in our networks. However, Ising couplings are symmetric and do not abide Dale’s law. Hence, our study does not provide true insights into the role played by inhibition – an issue that shall be examined in future work. Along similar lines, two-photon imaging experiments in mice suggested that the decision circuit has a laminar organization.^68^ However, since we cannot identify cortical layers in our current data, we do not address this issue here. Lastly, while the Ising model offers numerous advantages, assuming that all functional connections are symmetric seems unlikely for a decision circuit. Thus, future studies should investigate alternative network models and examine asymmetries in neuronal couplings.

These issues notwithstanding, our results demonstrate the presence of a stable, structured, and reproducible circuit in OFC. Historically, network models for economic decisions have been built based on theoretical notions dissociated from any measure or estimate of connectivity.^20–26, 66, 69, 70^ In this perspective, our study provides a first empirical assessment for the structure of this neural circuit. While future work will undoubtedly refine our current assessment, empirical appraisals of the circuit connectivity, such as that provided here, will ultimately lay the foundations for a new generation of data-driven computational models.

## Methods

### Subjects and experimental procedures

All experimental procedures conformed to the NIH *Guide for the care and Use of Laboratory Animals* and were approved by the Institutional Animal Care and Use Committee (IACUC) at Washington University.

Surgical protocols and the choice task closely resembled those described in previous studies^15^. Briefly, three adult male rhesus monkeys (*Macaca mulatta*; D, 10.6 kg; E, 9.5 kg; F, 10.8 kg) participated in the experiments. After familiarization and chair training, we implanted a head post and an oval recording chamber (main axes 30 mm and 50 mm) under general anesthesia. The chamber was centered on stereotaxic coordinates (AP 30, ML 0) and allowed access to bilateral OFC with coronal penetrations. Penetrations were guided by structural MRI scans (1 mm sections) obtained before and after surgery.

During the experiments, monkeys sat in an electrically insulated enclosure in front of a computer monitor (57 cm distance), with the heads restrained. The gaze direction was monitored with an infra-red video camera (Eyelink, SR Research). The behavioral task was controlled through a custom software (Monkeylogic) written in Matlab (MathWorks)^71^. Monkeys performed a standard juice-choice task. In each session, they chose between two juices labeled A and B (with A preferred) offered in variable amounts. The quantities offered in any given trial are indicated as *q_A_* and *q_B_*. Across trials, *q_A_* and *q_B_* varied in the ranges [0, *Q_A_*] and [0, *Q_B_*], respectively. At the beginning of the trial, the animal maintained central fixation for 1.5 s (fixation window: 2°). Two offers were presented simultaneously on the two sides of the fixation point (distance from fixation point: 7° of visual angle). Each offer was represented by a set of color squares; for each set, the color indicated the juice flavor, and the number of squares indicated the juice quantity (each square represented one quantum of juice). Offers were on display for a randomly variable delay (1-2 s; uniform distribution), at the end of which the central fixation point was extinguished (go signal). The animal indicated its choice with a saccade and maintained peripheral fixation for an additional 0.75 s, after which the chosen juice was delivered. Different pairs of offers were presented pseudo-randomly. For each offer pair, the spatial configuration (left/right) was counter-balanced across trials. Sessions included 181-402 trials (median = 264 trials) and lasted 30-60 minutes. Across sessions and across monkeys, we used 12 different juice types, and 19 different juice pairings. The juice quantum was set at 60-100 μl in different sessions and was fixed within each session. Juice rewards were delivered via a custom-designed 3-way juice line. The present data set includes a total of 85 sessions (21, 39, and 25 sessions for monkeys D, E, and F, respectively).

Recordings focused on the orbital gyrus (area 13m). Neuronal data were collected from both hemispheres of monkeys D and E (left hemisphere, A 31:36, L −6:−9; right hemisphere, A 31:36, L 6:9) and from the right hemisphere of monkey F (A 30:36, L 6:10). Extra-cellular recordings were conducted using 32-channel linear arrays (V-probe, Plexon; 120 μm diameter), with contacts spaced 75 μm from each other. In each session, we used between 1 and 4 probes, with at most two probes in each hemisphere. Each probe was advanced using a custom-built motorized micro-drive with a step size of 1 μm. Electrical signals were amplified, band-pass filtered (100-7,500 Hz), and recorded at 40 kHz using a Plexon Omniplex system. Spike identification and sorting were performed using the Plexon Offline Sorter. Subsequent analyses were conducted in Matlab (version 2025a; MathWorks), unless otherwise noted.

Possible errors in spike sorting were mitigated in two ways. First, to avoid having a neuron recorded from multiple channels counted as separate cells, we excluded one neuron from any pair of cells exhibiting ≥10% of coincident spikes (i.e., spikes occurring within 1 ms from each other). Second, to avoid errors due to mistakenly splitting a single neuron into multiple units, we excluded pairs of cells recorded from the same electrode (i.e., at distance = 0) in all population analyses following network inference.

### Behavioral analysis and cell classification

For each session, choices were analyzed with a logistic regression.^72^ We used a log-value-ratio logit model defined as:

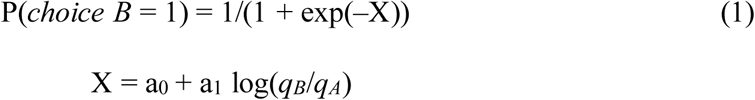

where *choice B* = 1 if B was chosen and 0 otherwise, and *q_A_* and *q_B_* were the juice quantities offered to the monkey. From the fitted parameters, we derived measures for the relative value *ρ* = exp(–a_0_ /a_1_) and the choice accuracy *η* = a_1_. In essence, the relative value *ρ* was the quantity ratio *q_B_*/*q_A_* that made the animal indifferent between the two juices and corresponded to the flex of the fitted sigmoid. The choice accuracy *η*, also termed inverse temperature, corresponded to the steepness of the fitted sigmoid.^72^

Previous studies indicated that different neurons in OFC encode variables *offer value A*, *offer value B*, *chosen juice*, and *chosen value*, and that each of these variables can be encoded with a positive or negative sign.^15, 46^ Thus in the analysis of individual neurons, we classified each cell as encoding one of these signed variables using the same procedures adopted in previous studies. Specifically, we defined 8 time windows aligned with different behavioral events: pre-offer (0.5 s preceding the offer onset), post-offer (0.5 s after the offer onset), late delay (0.5–1.0 s after the offer onset), pre-go (0.5 s preceding the go cue), reaction time (from the go cue to the saccade start), pre-juice (0.5 s preceding juice delivery), post-juice (0.5 s after juice delivery), and post-juice2 (0.5–1.0 s after juice delivery). An offer type was defined by two offered quantities; a trial type was defined by an offer type and a choice. For each cell and each time window, we averaged spike counts across trials for each trial type. A neuronal response was defined as the firing rate of one cell in one time window as a function of the trial type. Trial types with ≤4 trials were discarded from the analysis.

Cell classification proceeded in two main steps. First, for each neuronal response, we performed a Kruskal-Wallis test (factor = trial type). We imposed a significance threshold of p < 0.01. Neurons that satisfied this criterion in ≥1 time window were identified as task-related and included in subsequent analyses. Second, we defined variables

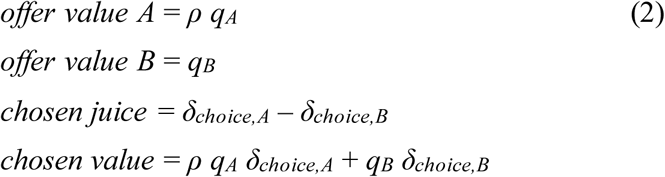

where *δ_choice,J_* = 1 if juice *J* was chosen and 0 otherwise. Each response passing the Kruskal-Wallis test was regressed on each variable. A variable was considered to explain the response if the regression slope differed significantly from zero (p < 0.05). If the variable explained the response, the sign of the regression slope defined the sign of the encoding (+ or –). Variables *chosen juice+* and *chosen juice*– were referred to as *chosen juice A* and *chosen juice B*, respectively. For each regression, we obtained the R^2^; for variables that did not explain the responses, we arbitrarily set R^2^ = 0. For each of the 8 signed variables (*offer value A+*, *offer value A*–, etc.), we summed the signed R^2^ across all time windows, and we obtained R^2^ .

Finally, each neuron was assigned to the signed variable that provided the maximum │R^2^│, where |·| indicates the absolute value. Neurons that did not pass the Kruskal-Wallis test in any time window or that were not explained by any variable were termed *unclassified*. As a result of this procedure, neurons were divided into 9 groups: *offer value A+*, *offer value A*–, *offer value B+*, *offer value B*–, *chosen juice A*, *chosen juice B*, *chosen value+*, *chosen value*–, and *unclassified*.

### Network inference

For each session, we fitted the activity of the neuronal population with an Ising model, which provided estimates for the functional connectivity between cell pairs. Spike times were examined in 5-ms bins, with a 1-ms sliding lag. The bin width was chosen to be sufficiently long to capture pairwise interactions (i.e., longer than synaptic delays), but sufficiently short so that neurons rarely spiked more than once per bin.

In any time bin, the activity of the N neurons recorded in the session was represented by a vector of binary variables, ***σ*** = (*σ*_1_, *σ*_2_, …, *σ_N_*), with *σ_i_* = 1 if cell *i* emitted a spike and 0 otherwise. For each trial, we analyzed the time bins from trial onset to trial end, excluding intertrial intervals.

For each neuron, time bins were concatenated across trials in the session. In the Ising model, the probability of observing a specific activity configuration is given by:

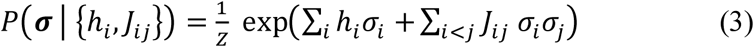

with:

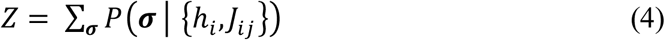

The partition function *Z* ensures normalization of the probability distribution in **Eq.3**. Parameters {ℎ*_i_*} represent external inputs that capture each cell’s intrinsic tendency to spike; couplings {*J_ij_*} capture the pairwise interactions between cells. Notably, the Ising model is the least constrained (maximum entropy) model consistent with given first- and second-order moments; as such, it captures pairwise interactions while excluding higher-order interactions.^35, 39, 40^ We define *f_i_* and *f_ij_* as the empirical mean firing rates and mean pairwise correlations – i.e., the average values over time bins of *σ_i_* and *σ_i_ σ_j_*, respectively. Fitting (or inferring) the Ising model means identifying the set of parameters {ℎ*_i_*, *J_ij_* } such that the average values of *σ_i_* (for all cells *i*) and *σ_i_ σ_j_* (for all pairs of cells *i*, *j*) under the probability distribution *P* match *f_i_* and *f_ij_*, respectively (**Eq.3**). Equivalent formulations of this problem include maximizing the log-likelihood of the empirical activity configurations under the model or minimizing the cross-entropy (i.e., the negative log-likelihood per time bin).

To ensure a unique and well-conditioned solution, we imposed a Gaussian (or L2) regularization, which penalizes large coupling values. The network inference problem is computationally hard because the time to compute *Z* increases exponentially with the number of neurons. Several approaches have been proposed^73–82^. Here we used the Adaptive Cluster Expansion (ACE) method ^32, 36, 45, 83^ (https://github.com/johnbarton/ACE), which was shown to outperform other procedures in both accuracy and computational efficiency. ACE solves the inference problem by computing an adaptive expansion of the regularized cross-entropy in terms of clusters of neurons, retaining only statistically significant contributions. This procedure reduces overfitting and substantially decreases computational cost.

To estimate statistical errors, we computed the Hessian (*χ*) of the regularized cross-entropy, also known as the Fisher information matrix. This step required computing the 2^nd^, 3^rd^, and 4^th^ order moments, and was implemented in C++ to handle the high computational load. Since the Hessian of the cross-entropy evaluated at the minimum characterizes its local curvature, the statistical uncertainty of each parameter can be estimated from the inverse of the Hessian matrix:

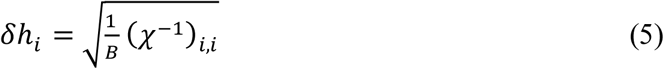

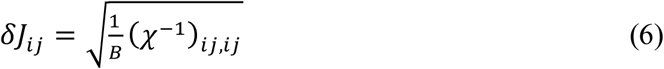

where B is the number of time bins in the session. A coupling *J_ij_* was considered statistically different from zero if its absolute value was at least 1.5 times its statistical uncertainty (|*J_ij_*| ≥ 1.5 *δJ_ij_*).

To assess the robustness of inferred networks, we performed 3 cross-validation procedures. First, we split spike data from each session into two interleaved sets of time bins (odd bins, even bins). We then inferred a separate network for each set, and we compared the resulting couplings. Second, we inferred a separate network in each of the 8 time windows defined above (pre-offer, post-offer, etc.). We then examined whether couplings were stable across the time windows. Third, we split trials into easy and hard decisions according to the decision difficulty, which we defined as follows:

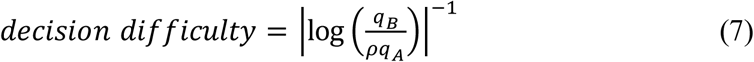

We inferred a network for each group of trials, and we compared the couplings.

### Predicting spike timing

The Ising model expresses the probability distribution of activity configurations – i.e., all possible patterns of active and silent neuronal states within a time bin – as a function of the fitted parameters ℎ and *J* (**Methods**, **Eq.3**) From this distribution, we can compute the conditional average activity of each neuron given the activity of all other cells. In other words, the inferred network provides a prediction for the probability that any given neuron spikes in a particular time bin, conditioned on which other cells spike in that time bin. We sought to assess the average accuracy of this prediction, and to compare this accuracy with that based on two “mean-field” predictions derived (1) from the average activity of the same neuron in other trials of the same type, and (2) from the average activity of other neurons of the same type in the same trial.

For each cell *i*, trial *r*, and time bin *t* we computed three spiking probabilities. Under the Ising model, the predicted spiking probability of cell *i* given the activity of the other cells in the same time bin *t* – indicated by ***σ***_−*i*,*r*,*t*_ – is a logistic function of the total input *I* that neuron *i* receives from the population:

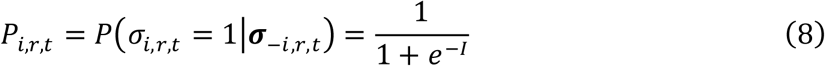

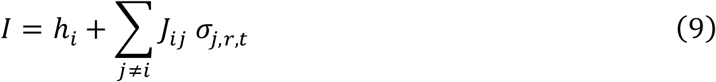

Under the mean-field model in which the cell’s spiking probability equals the average activity of the same cell in other trials of type *R* in the same time bin *t*, the predicted spiking probability is:

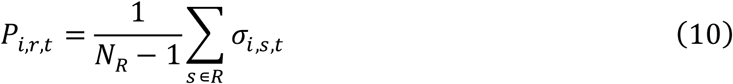

where *N_R_* is the number of trials of type *R*. Finally, under the mean-field model in which the cell’s spiking probability equals the average activity of the other cells of the same cell group *G* in the same time bin *t*, the predicted spiking probability is:

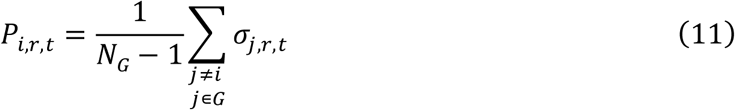

where *N_G_* is the number of cells in group *G*.

To compare the predictive power of the 3 models, we first binned the predicted spiking probabilities (*P_i_*_,*r*,*t*_) into 20 equally spaced probability bins spanning from 0 to 1. Notably, different models have different sets of time bins contributing to each probability bin. Then, for each model *m* and each probability bin *k*, we calculated the Spike Prediction Error (*SPE^m^*). This error was defined as the absolute difference between the actual spike frequency (*ASF^m^*) observed experimentally in the set of time bins associated with probability bin *k* for model *m* and the predicted spike probability (*PSP^m^_k_*) generated by the model:

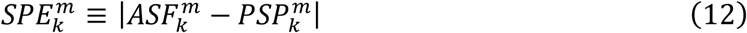

Finally, to compute a single, comprehensive error value for each model, we calculated the mean Spike Prediction Error (*SPE^m^*). This metric was a weighted average of the errors across probability bins, where the weight assigned to bin *k* was proportional to the base-10 logarithm of the number of time bins (*μ^m^_k_*) that contributed to that probability bin:

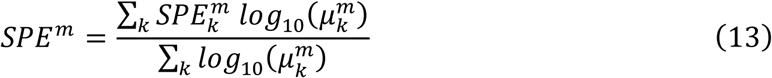

Alternative procedures to compute *SPE^m^* did not significantly alter the results. In particular, the results obtained replacing *log*_10_ (*μ_k_^m^*) with 1 or with *μ_k_^m^* were nearly identical to those illustrated in **Suppl. Fig.5**.

### Inferred couplings and correlation indices

A general question for network inference analyses is how the inferred connectivity compares to simple pairwise correlation^36, 47^. In fact, Ising couplings are most directly comparable to the correlation index, which is closely related to noise correlation. Specifically, given two neurons *i* and *j*, we defined the correlation index *CI_ij_* as:

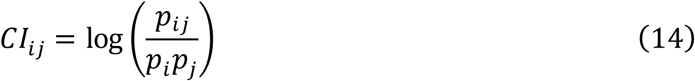

where *p_i_* is the probability that neuron *i* emits a spike in any given 5-ms time bin and *p_ij_* is the probability that both neurons *i* and *j* emit a spike in the same 5-ms time bin. For a system of only two neurons, assuming that *p_ij_* ≪ *p_i_* ≪ 1 (which is typically the case with 5-ms bins), it can be shown that *J_ij_* ≈ *CI_ij_*^36^. In larger networks, *J_ij_* and *CI_ij_* are in general not identical, but they are correlated.

To quantify the indirect connectivity beyond the second order, we defined an indirect coupling strength, *J^indirect^_ij_*, representing the net interaction between neuron *i* and neuron *j* mediated through third neurons *k* recorded simultaneously. This was calculated as the sum of the products of the component direct couplings for all possible intermediary neurons:

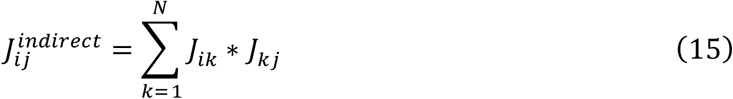

This metric characterized the prevalence and strength of specific indirect triplet coupling motifs within the recorded network.

### Mean coupling strength

Starting from couplings *J* inferred with the Ising model, we summarized interactions between functional cell groups by computing the mean coupling strength (*MCS*) – a 9×9 matrix where each row and each column represented a signed variable (a functional cell group). For each pair of cell groups *u* and *v*, entry *MCS_uv_* was defined as:

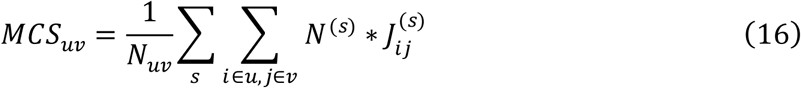

where *N_uv_* is the total number of simultaneously-recorded cell pairs encoding variables (*u*, *v*) available in our data set (i.e., the number of elements in the sum), *N*^(*s*)^ is the number of cells recorded in session *s*, and *J_ij_*^(*s*)^ is the inferred coupling between cells *i* and *j* recorded in session *s*. The sum is over all pairs of cells *i*, *j* such that *i* encodes variable *u* and *j* encodes variable *v*, and over all sessions. *MCS_uv_* captures the mean interaction between cell groups *u* and *v*, not that between a single cell pair.

Of note, the factor *N*^(*s*)^ in **Eq.16** weighted each inferred coupling by the number of cells recorded in that session. This correction was introduced because the mean absolute value of inferred couplings, mean(|*J_ij_*^(*s*)^ |), is expected to decrease as a function of the number of cells recorded simultaneously – a prediction confirmed in our data (**Suppl. Fig.6**). The correction ensured that estimates of *MCS* did not depend on the number of cells typically recorded in our experiments (i.e., on the mean 〈*N*^(*s*)^〉*_s_*), or on the session-to-session variability in that number.

### Defining and computing the effective network

The primary goal of this study was to analyze the architecture of the decision circuit at the mesoscale. To do so, we condensed the neuron-to-neuron connectivity inferred with the Ising model into a 9×9 effective network (*EN*) where each row and each column represented a signed variable (a functional cell group). This coarse-graining operation was conducted using a mean-field approach.

A reduced network between functional cell groups should preserve how activity in one group statistically influences the activity of another group. In the Ising model, that influence is captured by the conditional spiking probability:

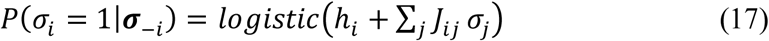

The quantity ℎ*_i_* + ∑*_j_ J_ij_ σ_j_* is the effective input acting on neuron *i*. Contribution ∑*_j_ J_ij_ σ_j_* determines how the activity of the rest of the network biases neuron *i* toward spiking or silence. Let us now consider the influence of cell group *v* on cell group *u*. If the activity of neurons in group *v* is statistically summarized by the mean activity *m_v_* = 〈*σ_j_*〉*_j_*_∈ *v*_, the average input received by each neuron *i* in group *u* from neurons in group *v* (in session *s*) is:

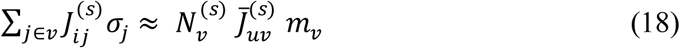

where *N*^(*s*)^_*v*_ is the number of neurons in group *v* recorded in session *s*,

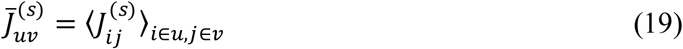

The effective group-to-group coupling for session *s* is defined as:

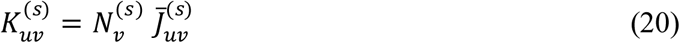

such that:

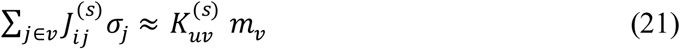

for every neuron *i* ∈ *u*. Reduced self couplings (diagonal entries) are computed as follows:

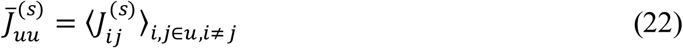

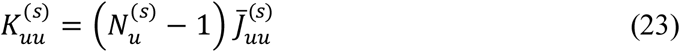

The effective coupling matrix *K*^(*s*)^ quantifies the increase mean activity of population *u* conditioned on the increase in mean activity of population *v* (in session *s*). Thus, it represents the effective statistical influence between functional populations. Notably, *K*^(*s*)^ is not symmetric.

The effective network (*EN*) was defined by averaging *K*^(*s*)^ across sessions:

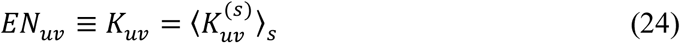

If the number of cells recorded in each session (namely *N*^(*s*)^) is sufficiently large, *EN* can be approximated as follows:

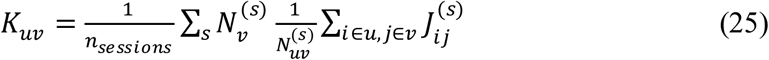

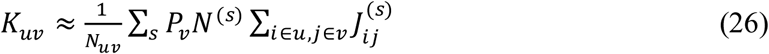

where *n_sessions_* = 85 is the number of sessions in our data set, *N* is the number of cell pairs (*i* ∈ *u*, *j* ∈ *v*) recorded in session *s*, and *P_v_* is the probability that a neuron encodes variable *v* (obtained from **Suppl. Table 3**). Finally, combining **Eq.26** and **Eq.16**, we obtain:

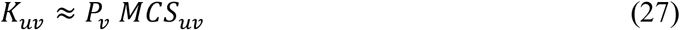

Using **Eq.27**, *EN* entries can be computed pooling cell pairs from the whole data set, as opposed to doing so for individual sessions first. This procedure made it possible to conduct the analyses described in the following sections.

### Assessing the reproducibility of the effective network and the mean coupling strengths

To assess reproducibility across monkeys, we computed *EN* separately for each animal, and we represented it as a 81-dimensional vector. For each pair of monkeys (D:E, D:F, E:F), we then calculated the Spearman correlation between the two vectors. We used a similar procedure to assess whether the *EN* was reproducible across hemispheres. In this case, we pooled data from different monkeys, separately for the left and right hemispheres. Similarly, we computed the reproducibility of *MCS*. However, since it was symmetric, *MCS* was represented as a 45-dimensional vector.

We examined how the structure of *EN* and *MCS* depended on the distance between cell pairs (**Fig.6A**). First, we approximated the position of each cell with that of the electrode it was recorded from. Then, we considered the distribution of cell-pair distances in our data set.

Focusing on cell pairs at distance ≤1 mm, we divided the distribution in 4 quantiles, each with the same number of cell pairs (N ≈ 15,000). Cell pairs at larger distances were divided into 3 additional pools: same probe (distance 1–2.325 mm); different probe, same hemisphere (distance 2.325–10 mm); different hemispheres. Thus we obtained 7 pools total. For each pool k = 1…7, we computed the *MCS* matrix, which we labeled *MCS*^(*k*)^. Then we plotted entries of *MCS*^(*k*)^ against the corresponding entries of *MCS*^(1)^, we computed the Spearman correlation, we performed a Deming linear regression, and we quantified the regression slope. Finally, we studied how the Spearman correlation and the regression slope varied as a function of k – i.e., as a function of the distance between cells. For pool 1, y-axis measures in **Fig.6A** were derived from a split-half procedure. Pools at larger distances (>1 mm) included larger numbers of cell pairs. Thus to isolate the effects of cell distance from those of sample size, we estimated *EN* and *MCS* for these pools using a bootstrap procedure. For each pool, we sampled with replacement the same number of pairs included in the first 4 pools; we then computed *EN* and *MCS*, and compared them to those obtained for k = 1. The procedure was repeated 1,000 times; reported values and error bars are bootstrap means and standard deviations.

We also examined the network reproducibility as a function of the number of cell pairs, separately for different pools of cell pairs defined by distance or treatment (**Fig.6B**). For each pool, we randomly selected N cell pairs, split them into two sets, computed the two *EN* matrices, and then computed the Spearman correlation between them. We repeated this procedure 100 times and computed the mean Spearman correlation (shown in the figure). For the “random cell group” treatment, we randomly reassigned each cell to a variable with probabilities matching the empirical distribution of cells across groups in the whole neuronal population (**Suppl. Table 3**).

The signal-to-noise ratio (SNR) of coupling strength was defined as the ratio between coupling strength and statistical error of coupling:

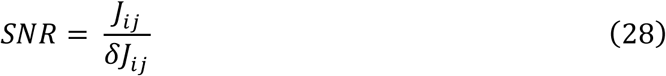

### Analysis of matching sessions

We identified pairs of matching sessions – i.e., sessions run with the same juice flavors and the same sets of offer values. We allowed ≤1 mismatch in trial types, and we imposed that the two relative values be very similar. Specifically, referring to the matching sessions as session 1 and session 2, we imposed |log(*ρ₁*/*ρ₂*)| < 0.1. In our data set, 78 pairs of sessions met these criteria. Because any given session could be matched with >1 session, some sessions contributed to multiple pairs. To verify that this duplication did not bias our results, we conducted a control analysis where we imposed that each session participate in at most one matching pair. With this restriction, our data set included 15 pairs of matching sessions.

For each pair of matching sessions, we matched individual trials and eliminated trials that could not be matched. Since offer period and reaction time varied from trial to trial, the duration of matched trials was generally unequal. We therefore defined two time windows aligned with the offer onset (from −500 ms to +1000 ms) and with the juice delivery onset (from −500 ms to +700 ms). We concatenated the two time windows for each trial, and then concatenated all trials within each session. This procedure yielded, for each of the two sessions, a data matrix where rows and columns were neurons and time bins, respectively. The two matrices had the same number of columns (because times were matched) but generally different numbers of rows (because cell counts could differ).

Given matching sessions 1 and 2, with N₁ and N₂ cells, respectively, we first performed the standard network inference for each session separately (*J^individual^*). Next, we combined the two data matrices by vertical concatenation, and we inferred the network for the combined pseudo-session. The resulting matrix of inferred couplings was partitioned into four blocks: two diagonal blocks (N₁×N₁ and N₂×N₂) containing couplings between pairs of cells recorded in the same session (*J**^ww^**^it^*^ℎ*in*^ *^session^*), and two off-diagonal blocks (N₁×N₂ and N2×N_1_) containing couplings between pairs of cells recorded in different sessions (*_J_^across^ ^sessions^*).

For each session pair *p* and each cell-group pair (*u*, *v*), we first computed a per-pair mean coupling for each treatment:

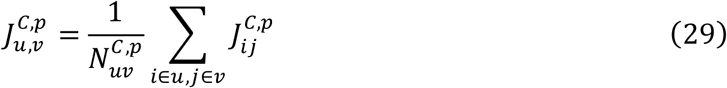

where *C* is the treatment (*individual*, *within session*, or *across sessions*) and *N^C^*^,*p*^ is the total number of cell pairs encoding variables (*u*, *v*) in session pair *p*, treatment *C*. The 9×9 matrix entry was then the unweighted average of these per-session pair means for all *P* session pairs:

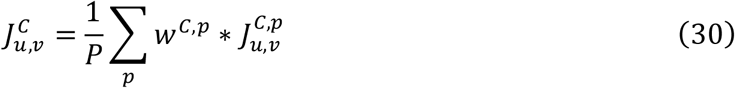

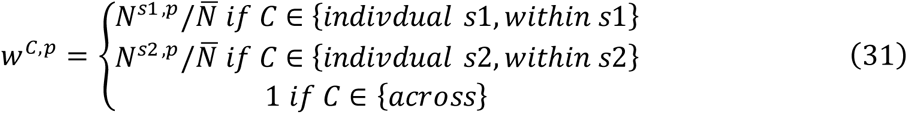

where *N^s^*^1,*p*^ and *N^s^*^2,*p*^ are the number of neurons in session 1 and session 2 of pair *p*, and *N̄* is the mean session size across all sessions included in pairs. Of note, where *w^C^*^,*p*^ depends on both the treatment *C* and the specific pair *p*.

Of note, this definition is nearly identical to that of *MCS* (**Eq.16**), except for the *N̄* normalization, which was introduced to estimate couplings at the scale of single cells. The *N*^(^*^s^*^)^/*N̄* factor was included for *individual* and *within session* treatments because the spiking probability of a cell *i* in session 1 is driven only by other neurons in session 1, not by neurons in session 2. No weight was included for the *across sessions* treatment because the spiking probability was only weakly driven by the behavioral variables and could not be scaled by the number of recorded cells.

In another analysis, we mismatched trials and misaligned time events for the two matching sessions. The *across sessions* couplings derived from this analysis are referred to as *J^mismatc^*^ℎ*ed*^. Specifically, we mismatched trials by pairing trials of session 1 randomly with trials of session 2. This procedure removed from *J^mismatc^*^ℎ*ed*^ any effect of trial variables *q_A_*, *q_B_*, and chosen juice.

Furthermore, for each trial of session 2, we randomly selected a time t0 within the 2,700 ms time window, and we wrapped times of all cells in the session around t0. In other words, we transformed the original time window [0 : 2,700] to the time window [[t0 : 2,700], [0 : t0-1]] (all times in ms). Because the same t0 was used for all cells in session 2, neurons recorded simultaneously remained temporally aligned, and within session statistics were preserved.

However, because sessions 1 and 2 had different temporal offsets, task-related inputs were mismatched across sessions. Thus, this procedure removed from *J^mismatc^*^ℎ*ed*^ any effect of task events.

### Relation between effective network and context variables

We examined how *MCS* depended on a variable capturing the internal state of the animal, namely the relative value *ρ*, and on a variable characterizing the behavioral context, namely quantity range ratio *Q_B_* ⁄*Q_A_*. For each variable, we examined the distribution across sessions and divided it into two sets (three sectors) based on the median distance from the center of the distribution (mean value). The two sets of sessions (near-center and far-from-center) had roughly the same number of sessions. The mean value of the variable under consideration (*ρ* or *Q_B_* ⁄*Q_A_*) was nearly the same for the two sets of sessions. However, the variable under consideration was nearly fixed across near-center sessions, and highly variable across far-from-center sessions. We then proceeded to compute the *MCS* matrices for the two sets of sessions. For each set, we considered the pool of inferred couplings, split it randomly in half, computed the two *MCS*s, and quantified their similarity (Spearman correlation). We repeated this procedure N = 1,000 times and obtained a bootstrap distribution.

Behavioral variables *ρ* and *Q_B_* ⁄*Q_A_* were partially correlated across sessions. To isolate the impact of each variable on the structure of *MCS*, we proceeded with a more granular analysis. Specifically, we examined the joint distribution of the two variables and we computed, for each variable, the median distance from the center of the marginal distribution. On this basis, we divided sessions in four sets with (a) fixed *Q_B_* ⁄*Q_A_*, fixed *ρ*; (b) fixed *Q_B_* ⁄*Q_A_*, variable *ρ*; (c) variable *Q_B_* ⁄*Q_A_*, fixed *ρ*; and (d) variable *Q_B_* ⁄*Q_A_*, variable *ρ*. We then repeated the split-test reliability test for each of sets (a), (b), and (c).

## Acknowledgements

We thank Timothy Kim and Alessandro Livi for comments on the manuscript. This work was supported by the NIH (grants R01-MH104494 and R01-DA032758 to CPS), the Sloan Foundation (Research Fellowship Award No. FG-2024-22163 to GT), the McDonnell Center for Systems Neuroscience (Small Grant GF0013150 to GT), the Japan Student Services Organization (predoctoral fellowship to YK), and Washington University (BioSURF Fellowship Award for undergraduate research to TGC).

## Contributions

CPS and GT designed the study; HS trained the animals; YK and CPS performed the experiments with assistance from HS; YK and CPS processed the data and performed single-cell analyses; GT and TGC developed an automated software package to run the network-inference algorithm on the data; YK and GT analyzed the inferred networks; YK, TGC, KZ, MK, GT and CPS discussed and evaluated the results; YK and CPS wrote the initial draft; all the authors edited the manuscript. GT and CPS co-directed the project.

---

**Supplementary Figure 1.**
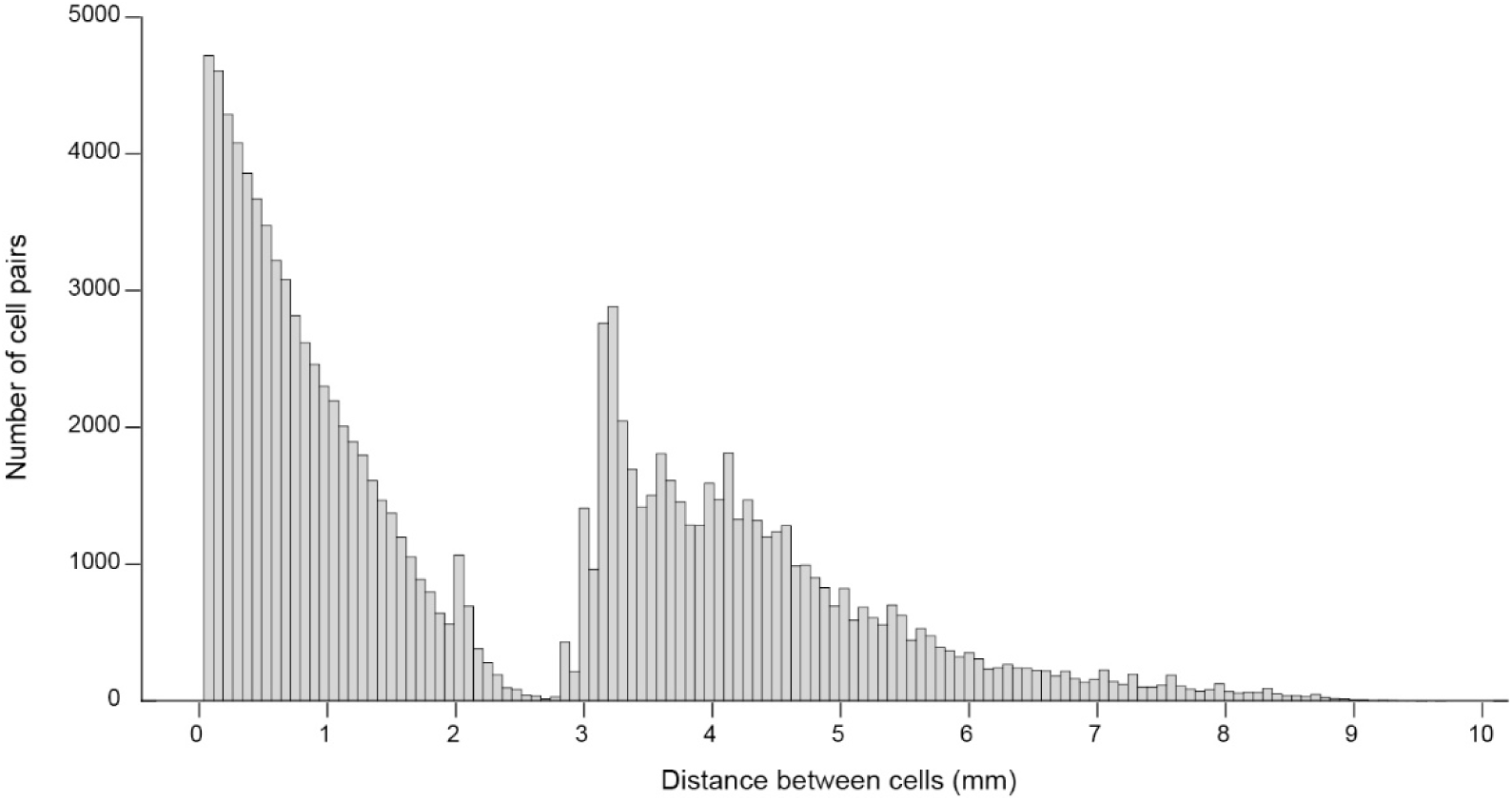
Distribution of cell pair distances. Distribution of distances between pairs of cells recorded in the same hemisphere. Data from 85 sessions (3 monkeys) were pooled. The data set included 118,028 cell pairs. For distances ≤ 2 mm, both cells were recorded from the same probe, and several sessions had probes placed to have cell pairs between 2–2.325 mm. Distance ≥2.325 mm implies that the two cells were recorded in the same hemisphere but from different probes. In addition, 30,997 pairs of cells were recorded in opposite hemispheres (not included in this figure).

**Supplementary Figure 2.**
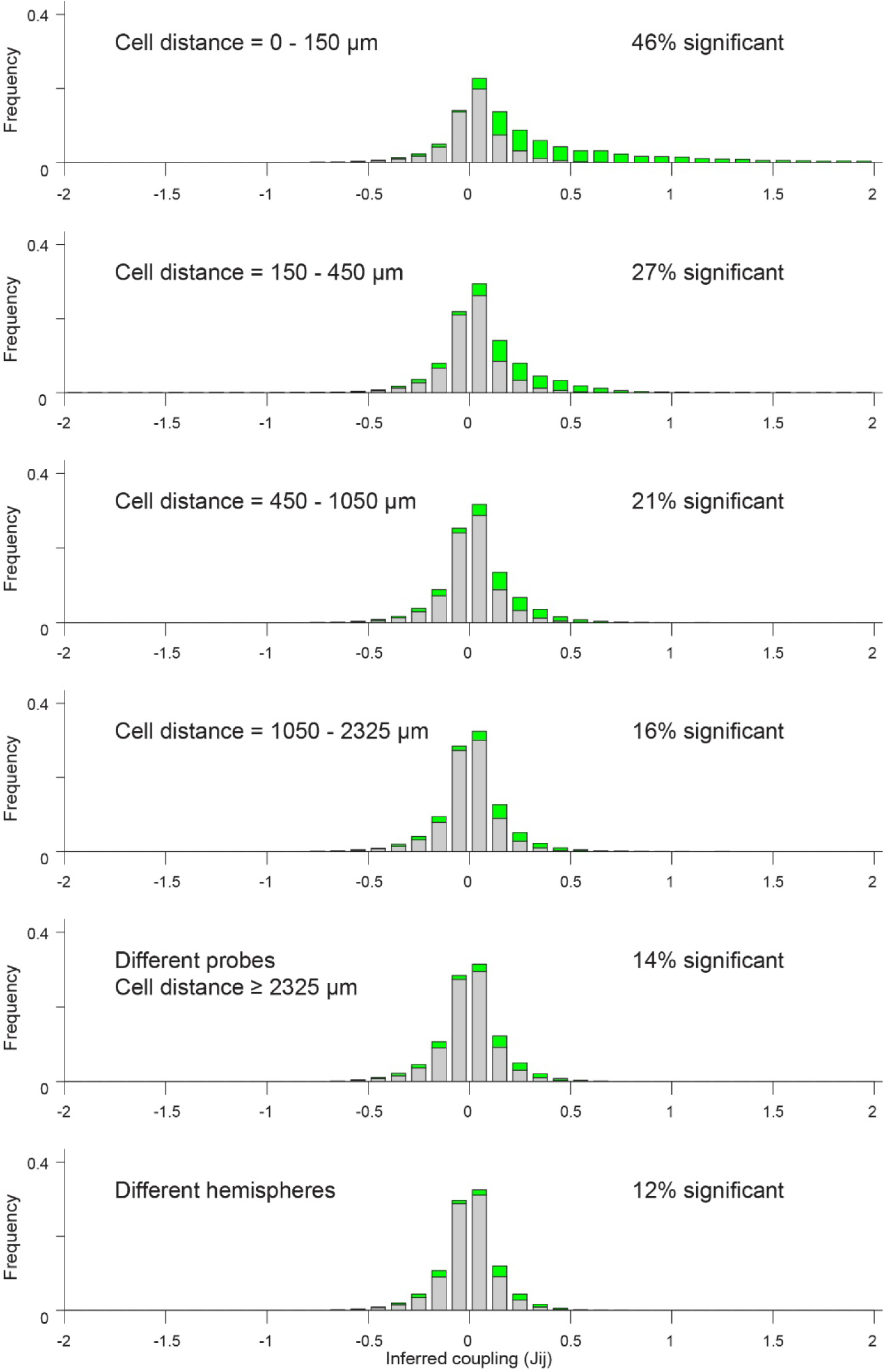
**Distribution of inferred couplings at different distances**. Distribution of inferred couplings (*J*) as a function of the distance between cells. Panels from top to bottom are for increasing distances, as indicated. Green and gray bars represent significant and non-significant couplings, respectively. The sum of frequencies for gray and green was normalized to 1 for each histogram. The proportion of significant couplings (indicated in each panel) decreased with the distance between cells.

**Supplementary Figure 3.**
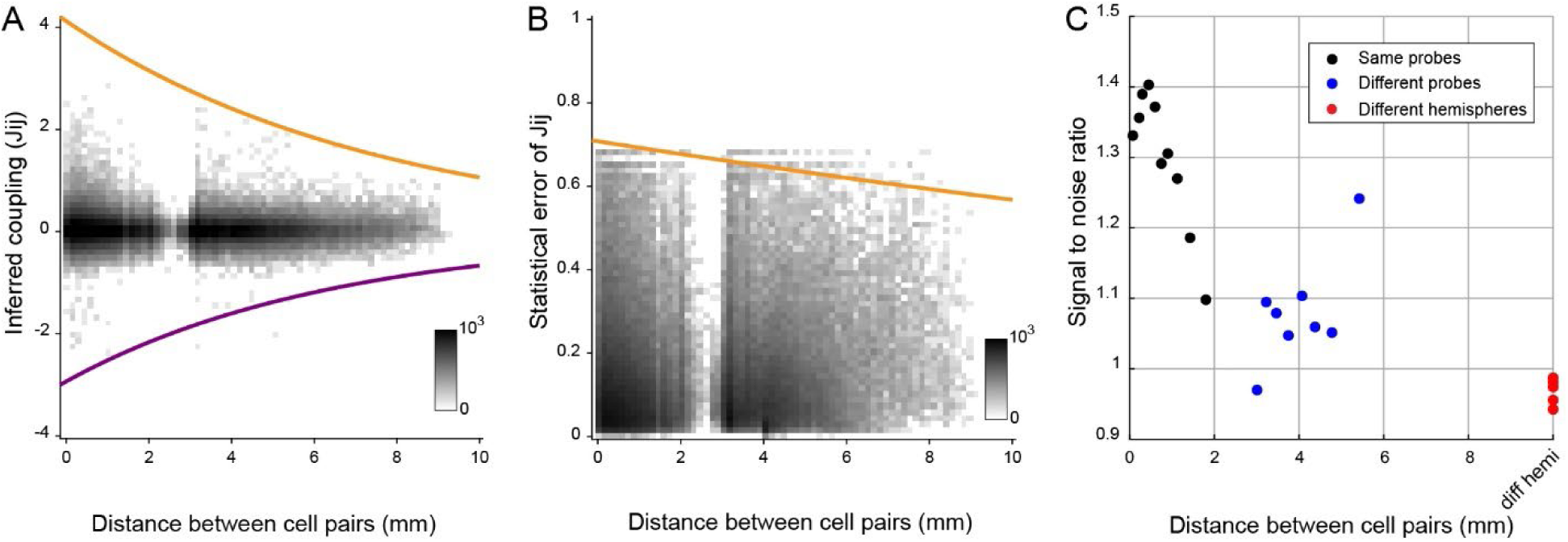
Signal-to-noise ratio of inferred couplings as a function of the cell distance. **A.** Inferred couplings as a function of the distance between cells. Because of the large number of data points, we display a pixel average of cell pair counts (pixel size = 0.05 mm x 0.14). Orange and purple curves were obtained by dividing distances in 30 bins of equal size, by computing the maximum absolute value within each bin, and by fitting these maxima with an exponential function. **B**. Statistical error of inferred couplings as a function of the distance between cells. We display a pixel average (pixel size = 0.05 mm × 0.014). Orange curve was obtained by the maximum value of the statistical error in each distance bin. **C**. Signal-to-noise ratio (SNR) of coupling strength as a function of the distance between cells. Cell pairs are sorted by distance into quantiles of 6,000 cell pairs. Each quantile is plotted as a point whose x-axis is the mean distance within the quantile and y-axis is the mean per-pair statistical SNR defined in **Eq.28**. Colors indicate cell pairs from the same probe (black), from different probes in the same hemisphere (blue), and from different hemispheres (red).

**Supplementary Figure 4.**
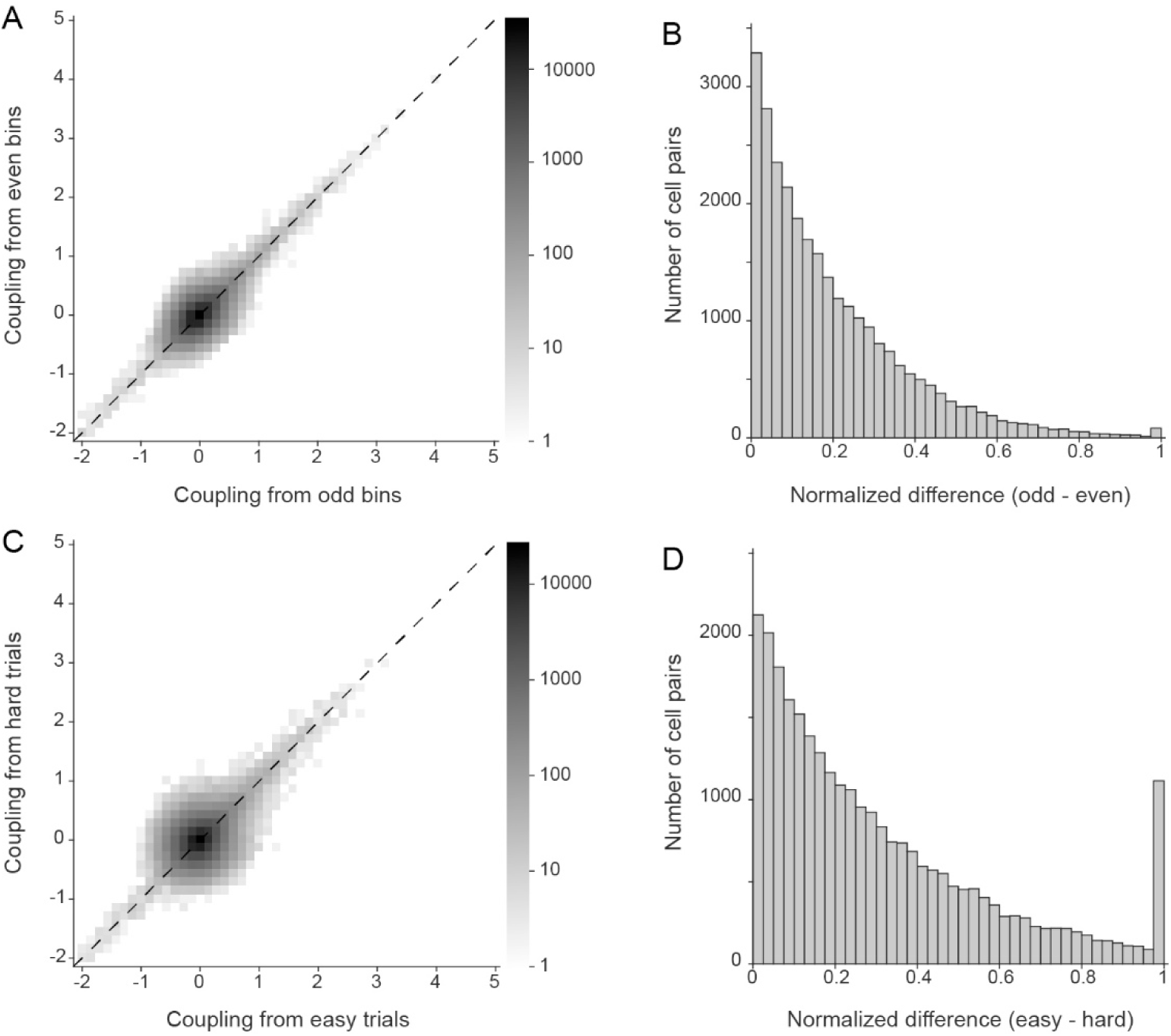
Inferred couplings are reliable and robust. **A**. Comparing couplings inferred in temporally subsampled data sets. In the scatter plot, x- and y-axis are couplings inferred from odd time bins and even time bins, respectively. Each cell pair contributed one data point. Because of the large number of data points, we display a pixel average (pixel size = 0.14 x 0.14). Couplings inferred in the two sets were highly correlated (corr = 0.49, 0.90 for statistically significant pairs). **B**. Distribution of normalized differences (odds vs. even time bins). The normalized difference was defined as |*J^odd^* − *J^even^*|⁄(|*J^odd^*| + |*J^even^* |) and computed for each cell pair. **C**. Comparing couplings inferred on easy vs hard decisions. In each session, we divided trials in easy and hard decisions (see **Methods**), and we inferred a network for each group of trials. Couplings inferred for the two groups of trials were highly correlated (corr = 0.24 for all cell pairs, 0.80 for statistically significant cell pairs). **D**. Distribution of normalized differences (easy vs hard decisions).

**Supplementary Figure 5.**
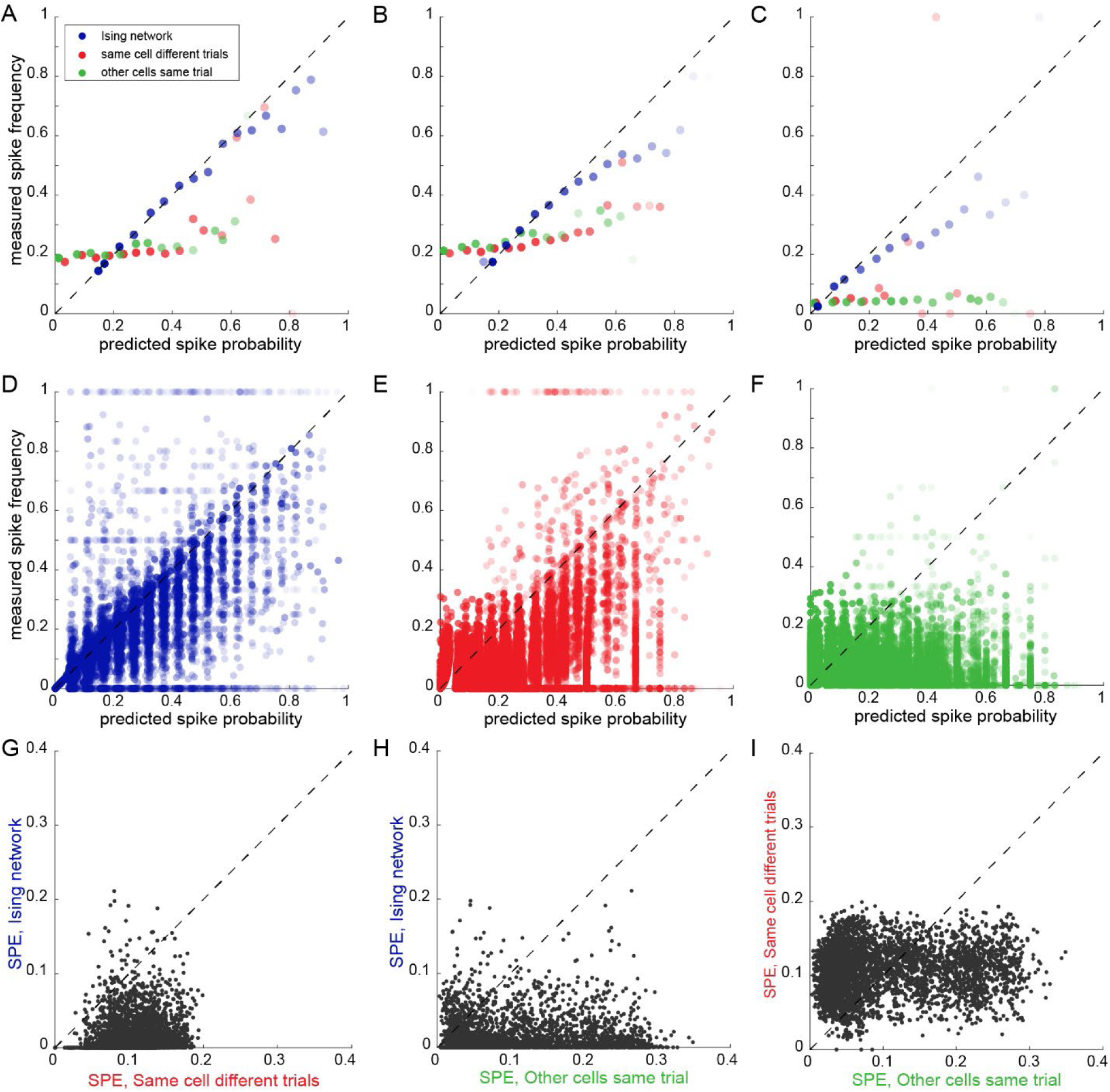
Predicting spike timing. For each neuron, we computed the spike probability in each 5-ms time bin based on the inferred Ising model (see **Methods**). The resulting distribution of probabilities across time bins was generally skewed towards zero because firing rates in OFC are relatively low. However, in some time bins, the probability could be substantially larger than zero. For each 5-ms time bin, we also knew whether the cell had actually emitted a spike. To test the accuracy of predictions based on the Ising model, we divided the probability interval [0, 1] into 20 bins of equal width. For each probability bin, we examined the 5-ms time bins for which the predicted spike probability fell in the probability bin, and we computed the actual spike frequency (average number of spikes over time bins). Finally, we compared the actual spike frequency with the predicted spike probability. To gauge the accuracy of Ising predictions, for each cell and each time bin, we also computed the spiking probability using two mean-field procedures. In one procedure, the predicted activity of the cell in a trial was equal to the average activity of the same cell in other trials of the same type (same cell, other trials). In the other procedure, the predicted activity of the cell in a trial was equal to the mean activity of other cells of the same type in the same trial (other cells, same trial). Each of these mean-field models provided a spiking probability in each time bin. **A**. Example cell. The panel illustrates the mean spike frequency (y-axis) as a function of the expected spike probability (x-axis), separately for the three models (see color legend). Each symbol represents one probability bin, and the opacity captures the number of time bins contributing to the data point. The dashed identity line corresponds to a perfect prediction. Notably, the Ising model provided a much better prediction for the mean spike frequency compared to the two mean-field models. **BC.** Two other example cells. **DEF.** Population analysis. We repeated the analysis illustrated in panels A-C for every neuron, and we pooled the results from the entire data set. Panel D illustrates the relation between mean spike frequency and expected spike probability predicted with the inferred Ising model. Here, each data point represents the activity of one cell in one probability bin. The other panels illustrate the equivalent plots obtained for the two mean-field models (E: same cell, other trials; F: other cells, same trial). Notably, data points are closer to the identity line in panel D (Ising) than in the other panels. In other words, the Ising model provided better predictions. **GHI.** Analysis of spike prediction errors. To quantify this observation of panels DEF, for each model, each cell, and each probability bin, we computed the difference between the actual spike frequency and the predicted spike probability. For each model and each cell, the spike prediction error (*SPE*) was defined as the weighted average of the absolute value of this difference. The 3 scatter plots contrast the *SPE* obtained from different models, pairwise. Each data point represents one cell. Notably, the Ising model (y-axis in panels G and H) vastly outperformed the two mean-field models (lower *SPE*). In all panels, color conventions are as in panel A. Control analyses confirmed that these results did not depend on the precise definition of *SPE* (not shown).

**Supplementary Figure 6.**
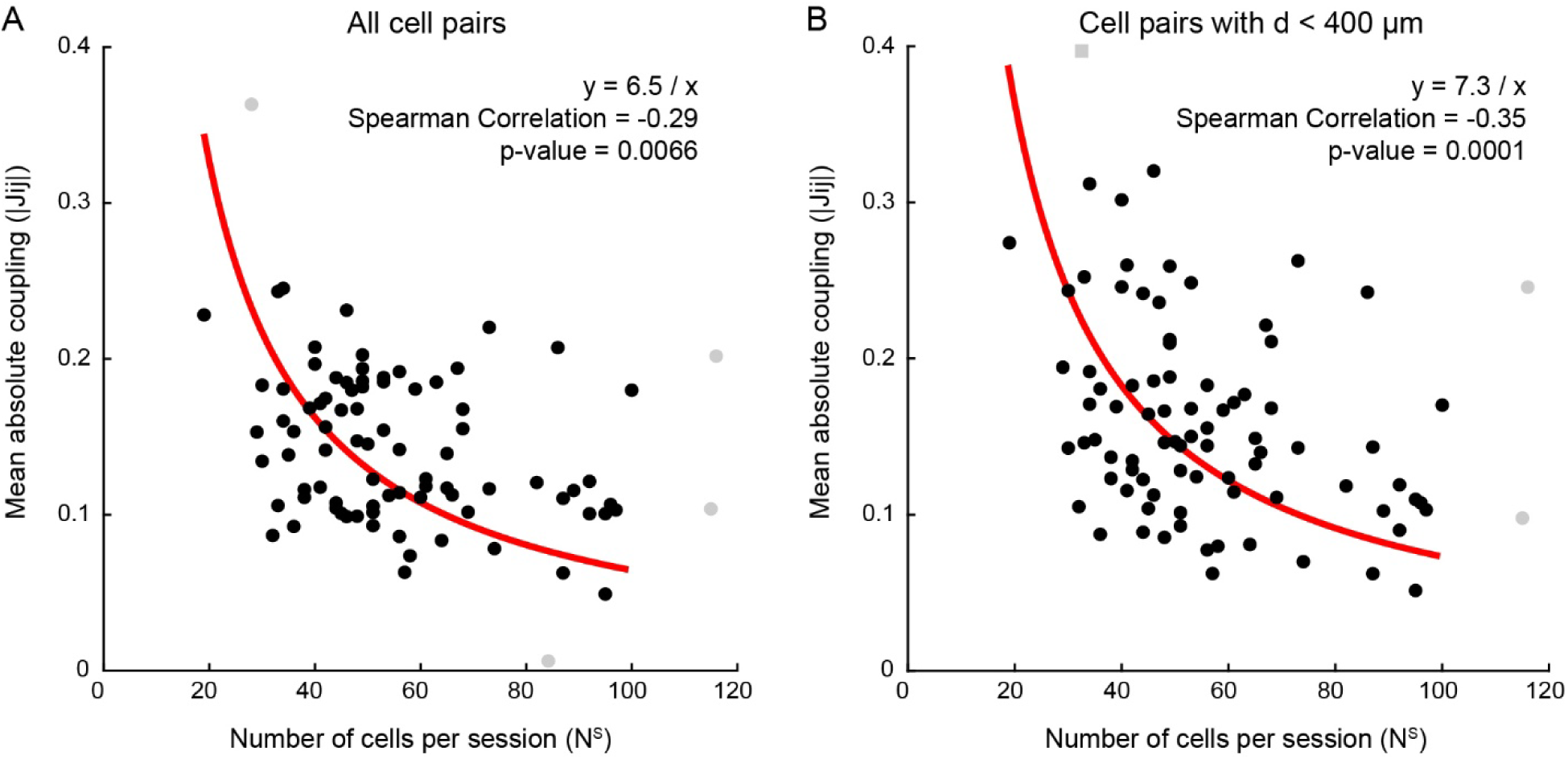
Mean inferred couplings decreased as a function of the number of recorded cells. **A.** The mean absolute value of the inferred couplings (*mean*(|*J_ij_* |); y-axis) is plotted here against the number of cells recorded in the session (*N^S^*). Each data point represents one session and all cell pairs are included. The two measures were significantly anti-correlated. The red line was obtained from a linear regression of the mean absolute coupling against 1/N^S^. Shaded points indicate outliers excluded from the calculation of the correlation. **B**. Analysis restricted to cell pairs at short distance. In our data set, in sessions with larger number of neurons, the mean distance between cell pairs was generally larger (e.g., because we recorded with more probes). Thus, in principle, the result of panel A could simply reflect the relation between *mean*(|*J_ij_*|) and cell distance (**Suppl.** Fig.3). To address this issue, we repeated the analysis including in the computation of *mean*(|*J_ij_*|) only pairs of cells at distance ≤400 μm, and replicated the effect. The inverse relation between *mean*(|*J_ij_*|) and *N^S^* arises because the same conditional spiking probability of a given cell *i* (**Eq. 8** and **Eq.9**) can be explained by contributions from a larger number of recorded cells *j* when *N^S^* is larger. As a result, the relative contribution of each cell *j* to the spiking probability of cell *i* is, on average, reduced (i.e., the absolute coupling |*J_ij_* | is smaller).

**Supplementary Figure 7.**
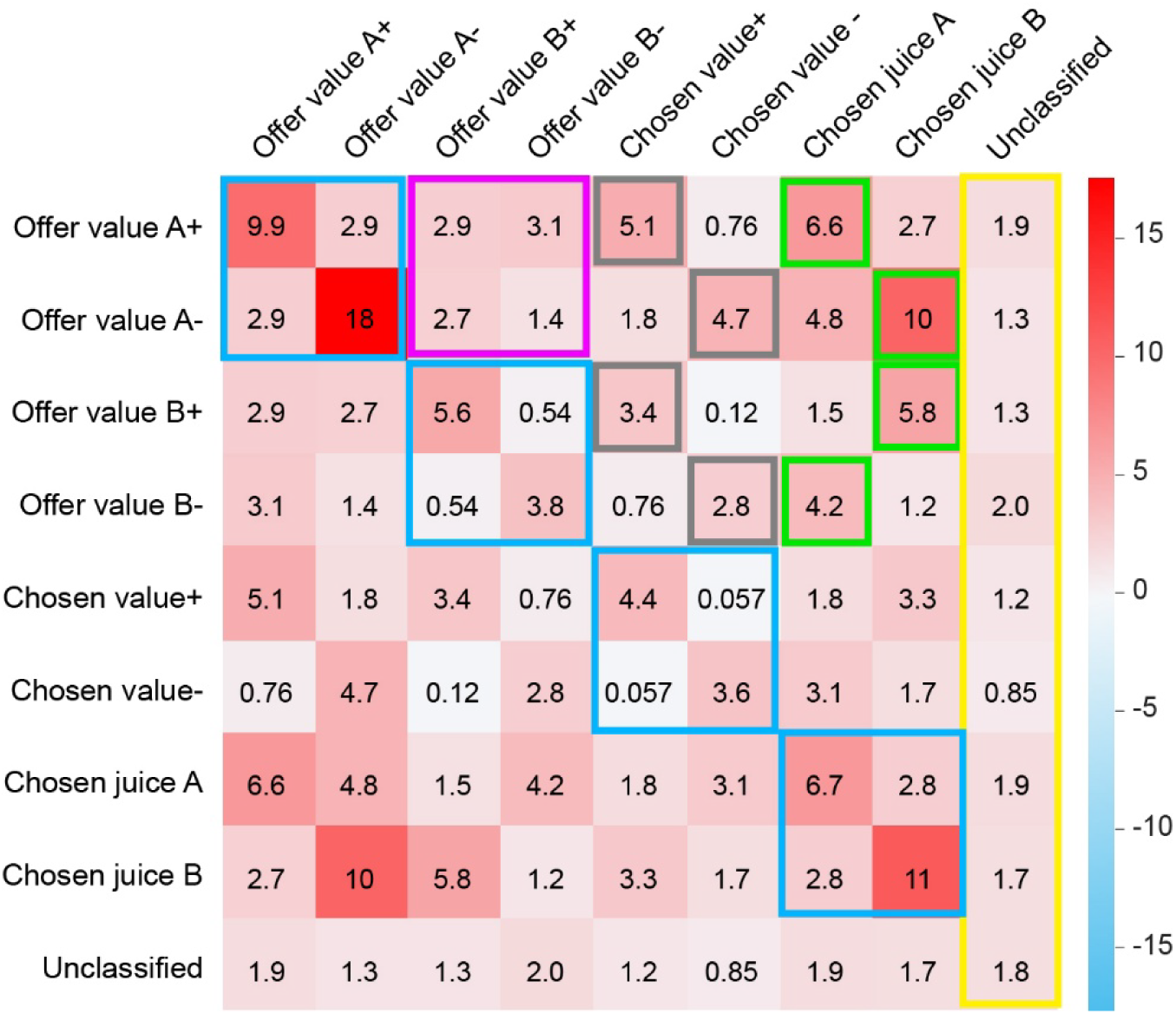
Structure of *MCS*. The matrix shown here is the same as Fig.3D, except that we highlighted the most notable traits. (1) Couplings between classified and unclassified cells were generally weak and unstructured (yellow). Their mean value could be taken as a baseline comparison for other entries of the matrix. (2) For each of the 4 variables (*offer value A*, *offer value B*, *chosen value*, *chosen juice*), couplings between cells encoding it with the same sign were enhanced, while couplings between cells encoding it with opposite signs were at or below baseline (blue). (3) Couplings between offer value cells and chosen juice cells were enhanced when the two neurons supported the same decision compared to when they supported opposite decisions (green). (4) Couplings between *offer value* cells associated with different juices were unstructured and barely above baseline (magenta). (5) Couplings between *offer value* cells and *chosen value* cells were enhanced when the encoding sign was the same and depressed when the encoding sign was opposite (gray).

**Supplementary Figure 8.**
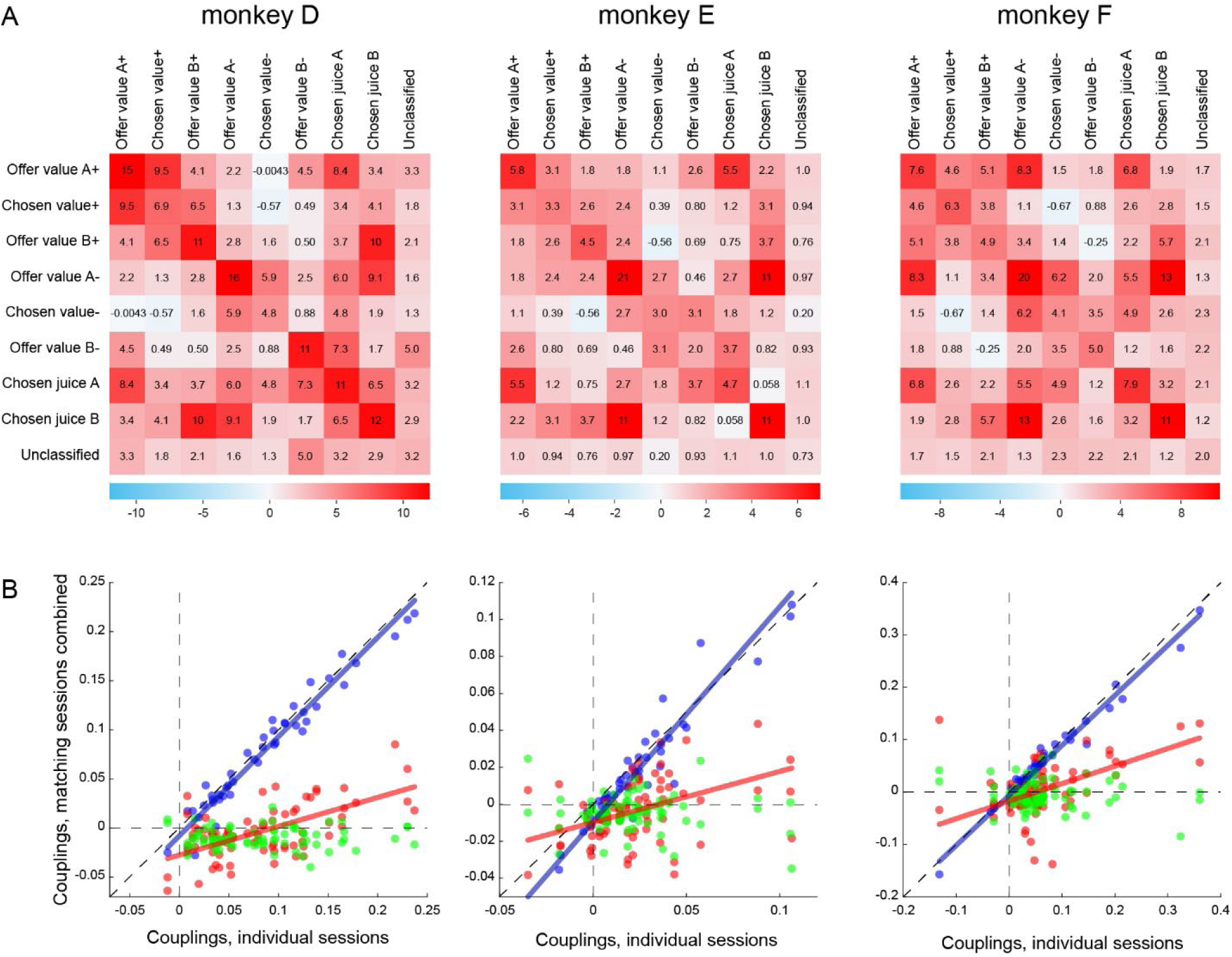
Analysis of individual animals. **A.** Mean coupling strength (*MCS*) computed for individual monkeys. Same conventions as in Fig.3D. **B.** Dependence on behavioral conditions. Same conventions as in Fig.4DE. Only regression lines for which the slope differed significantly from zero (p<0.05) are shown.

**Supplementary Table 1.** Number of cells and cell pairs by monkey.

|  | N sessions | N cells | N cell pairs |
| --- | --- | --- | --- |
| Monkey D | 21 | 1,260 | 43,249 |
| Monkey E | 39 | 2,417 | 80,672 |
| Monkey F | 25 | 1,107 | 25,104 |
| Total | 85 | 4,784 | 149,025 |

**Supplementary Table 2.** Number of cells and cell pairs by Hemisphere.

|  | N sessions | N cells | N cell pairs |
| --- | --- | --- | --- |
| Left Hemisphere | 42 | 1,073 | 21,057 |
| Right Hemisphere | 80 | 3,711 | 98,753 |

**Supplementary Table 3.** Number of cells by the encoded variable.

|  | N cells |
| --- | --- |
| *offer value A +* | 210 |
| *offer value A –* | 84 |
| *offer value B +* | 342 |
| *offer value B –* | 116 |
| *chosen juice A* | 279 |
| *chosen juice B* | 257 |
| *chosen value +* | 452 |
| *chosen value –* | 202 |
| *unclassified* | 2,842 |

**Supplementary Table 4.** Number of cell pairs by the pair of encoded variables.

|  | *offer value A +* | *offer value A –* | *offer value B +* | *offer value B –* | *chosen juice A* | *chosen juice B* | *chosen value +* | *chosen value –* | *unclassified* |
| --- | --- | --- | --- | --- | --- | --- | --- | --- | --- |
| *offer value A +* | 322 | 251 | 804 | 281 | 721 | 705 | 1,202 | 500 | 7,145 |
| *offer value A –* |  | 63 | 375 | 127 | 300 | 318 | 460 | 243 | 2,876 |
| *offer value B +* |  |  | 894 | 545 | 1,277 | 1,123 | 2,365 | 863 | 10,940 |
| *offer value B –* |  |  |  | 88 | 462 | 385 | 798 | 312 | 4,141 |
| *chosen juice A* |  |  |  |  | 642 | 967 | 1,717 | 685 | 9,229 |
| *chosen juice B* |  |  |  |  |  | 469 | 1,459 | 633 | 9,229 |
| *chosen value +* |  |  |  |  |  |  | 1,673 | 1,152 | 15,081 |
| *chosen value –* |  |  |  |  |  |  |  | 270 | 9,227 |
| *unclassified* |  |  |  |  |  |  |  |  | 57,969 |

